# Loss of function in the autism and schizophrenia-associated gene *CYFIP1* in human microglia supports a role in synaptic pruning

**DOI:** 10.1101/2022.10.24.513576

**Authors:** Steven D. Sheridan, Joy E. Horng, Hana Yeh, Liam McCrea, Ting Fu, Roy H. Perlis

## Abstract

**Background:** The *CYFIP1* gene, located in the neurodevelopmental risk locus 15q11.2, is highly expressed in microglia, but its role in human microglial function as it relates to neurodevelopment is not well understood.

**Methods:** We generated multiple CRISPR knockouts *of CYFIP1* in patient-derived models of microglia to characterize function and phenotype. Using microglia-like cells reprogrammed from peripheral blood mononuclear cells, we quantified phagocytosis of synaptosomes (isolated and purified synaptic vesicles) from human iPSC-derived neuronal cultures as an *in vitro* model of synaptic pruning. We repeated these analyses in human iPSC-derived microglia, and characterized microglial development and function through morphology and motility.

**Results:** *CYFIP1* knockout using orthogonal CRISPR constructs in multiple patient-derived cell lines was associated with statistically significant decrease in synaptic vesicle phagocytosis in microglia models derived from both PBMCs and iPSCs (p<0.0001). Morphology was also shifted toward a more ramified profile (p<0.0001), and motility was significantly reduced (p<0.0001). However, iPSC-*CYFIP1* knockout lines retained the ability to differentiate to functional microglia. *Conclusion*: The changes in microglial phenotype and function from loss of *CYFIP1* may contribute to pruning abnormalities observed in *CYFIP1*-associated neurodevelopmental disorders. Investigating risk genes in a range of CNS cell types may be required to fully understand the way in which common and rare variants intersect to yield neuropsychiatric disorders.

## Introduction

The 15q11.2 (BP1-BP2) microdeletion has been associated with an increased risk of multiple neurodevelopmental and neuropsychiatric disorders including epilepsy, autism spectrum disorders, intellectual disability and schizophrenia.(1-3) The presence of the 15q11.2 CNV increases the risk of abnormal neurodevelopment with a penetrance of as much as 10.4% (4), roughly doubling the risk of schizophrenia.(5) This locus contains four genes: non-imprinted in Prader-Willi/Angelman syndrome 1 gene (*NIPA1*), non-imprinted in Prader-Willi/Angelman syndrome 2 gene (*NIPA2*), cytoplasmic FMR1 interacting protein 1 (*CYFIP1*) and tubulin gamma complex associated protein 5 gene (*TUBGCP5*) (1).

Among these, *CYFIP1* has been most implicated in brain development, with recognized roles in synaptic functions including dendritic spine formation (6, 7), morphology (8) and branching.(6) In the neuronal context, *CYFIP1* interacts with the fragile X mental retardation protein (FMRP) to repress translation of neuronal genes, linking it to neurodevelopmental and neuropsychiatric disorders.(9)

Conversely, very little is understood about the non-neuronal functions of *CYFIP1* that may also contribute to central nervous system (CNS) development and dysfunction. In particular, the greatest levels of *CYFIP1* expression in CNS is in microglia, demonstrated by both mouse (10-12) and human (13) studies. However, there have been few studies of *CYFIP1* in these cells, particularly regarding possible mechanistic links to neurodevelopmental disorders. To further understand the potential roles of *CYFIP1* in such microglia processes, we used human *in vitro* models of synaptic pruning, in combination with Clustered Regularly Interspaced Short Palindromic Repeat (CRISPR) genome engineering. Although *CYFIP1* knockout in mice results in early embryonic lethality (6, 14), we showed that human induced pluripotent stem cells (iPSCs) lacking *CYFIP1* expression retained pluripotency upon iPSC clonal expansion, extended proliferation and the capacity to be differentiated into microglia. Using multiple orthogonal *in vitro* engineered human microglia models, including reprogrammed PBMC and iPSC-differentiated microglia, we demonstrated that loss of *CYFIP1* expression results in greatly reduced phagocytosis of human synaptic vesicles. Additionally, iPSC-differentiated microglia revealed altered morphology and motility upon further characterization, demonstrating additional roles of *CYFIP1* in microglia function. Taken together, these observations identified previously uncharacterized roles of *CYFIP1* that may contribute to aberrant microglial function in the developing CNS of patients with 15q11.2 CNVs or other variants in the *CYFIP1* gene that diminish function.

## Methods and Materials

### Preparation of CRISPR-Ribonucleoproteins (RNPs)

Single guide RNAs (sgRNAs, Table S1) were suspended to 50 uM in 1 mM Tris-EDTA, pH 8 and complexed with SpCas9 (IDT, #1081059) at a 1.5:1 sgRNA:Cas9 mole ratio for 10 minutes at room temperature. Two-part gRNAs comprising CRISPR-Cas9 crRNA (IDT) and tracrRNA (IDT, #1072533) (Table S1) were suspended to 100 uM in 1 mM Tris-EDTA pH 8, mixed at a 1:1 mole ratio, annealed by heating to 85C for 5 min, then cooling to 4C. Complexation with SpCas9 occurred at a 2:1 gRNA:Cas9 mol ratio.

### PBMC-derived microglia culture and transfection

Frozen and aliquoted peripheral blood mononuclear cells isolated from units of whole blood from healthy control subjects were purchased (Vitrologic, Inc. Charleston SC) and samples were quick-thawed in a 37 C water bath and immediately transferred into RPMI-1640 (Sigma, #R8758) + 10% heat-inactivated fetal bovine serum (Sigma, #12306C) + 1% of Penicillin/Streptomycin (Life Technologies, cat# 15140-122), washed by centrifugation at 300 g for 5 min followed by a media change and cell resuspension. They were plated at a density of approx. 30K cells/cm^2^ into a tissue culture treated vessel pre-coated for 1 hr at 37 C with Geltrex (Gibco, #A1413202) and cultured at 37C + 5% CO2. After 24 hrs, the media was carefully removed and replaced with RPMI-1640 + 1% Penicillin/Streptomycin + 1% Glutamax (Life Technologies, # 35050-061) + 100 ng/ml IL-34 (PeproTech, #200-34; Biolegend Inc, #577904) + 10 ng/ml GM-CSF (PeproTech, #300-03).

After an additional 8 days of differentiation, PBMC-derived microglia were harvested, saving the media, with Accutase (Sigma, #A6964) incubation at 37C until just unadhered, centrifuged at 300 g for 5 min and counted by hemocytometer. Using the 4D-Nucleofector X system (Lonza Biosciences, #AAF-1003X), all cells were resuspended in nucleofection solution, and 20,000 cells were mixed with each CRISPR-RNP, then transferred into cuvettes and subject to electroporation. The transfected cells were then moved into a well of a 96-well plate pre-filled with the original culture medium and cultured for an additional 6 days, until day 14.

### iPSC culture and transfection

Induced pluripotent stem cells (iPSCs) were generated from fibroblasts obtained from healthy control male donor lines PSC-01-179; 29-year old and PSC-01-092; 45-year old (Table S2) using synthetic mRNA pluripotency factors by Cellular Reprogramming, Inc. (https://www.cellular-reprogramming.com). They were cultured in 37C with 5% CO2 on geltrex-coated tissue culture plates in Essential 8 complete medium (E8; Gibco, #A1517001) with daily media changes. iPSCs were passaged as needed in the presence of 2 μM Rho-associated kinase (ROCK) inhibitor (RI) thiazovivin (Santa Cruz Biotechnology, #SC-361380). Prior to transfection, iPSCs were incubated in E8+RI for at least an hour. Transfections of CRISPR-RNPs to iPSCs were performed using the Lonza 4D-Nucleofector system following manufacturer’s protocols using 100,000 iPSCs per reaction. Following transfection, 50-100 iPSCs were plated per 6-well plate well in E8+RI for manual selection of resulting clonal colonies and expansion. Clonal purity was verified through Sanger sequencing of the genomic loci. Normal iPSC karyotypes were confirmed (Fig. S2) using Applied Biosystems KaryoStat™ microarray assay (ThermoFisher Scientific KaryoStat+ Service).

### Differentiation of iPSC-derived microglia (iPSC-iMGs)

Microglia were generated from clonal iPSCs adapted from a protocol that has been previously described.(15) In brief, iPSCs grown in E8 complete medium were transitioned into mTeSR Plus complete medium (StemCell Technologies, #100-0276) and MACs sorted for TRA-1-60+ cells (Miltenyi, #130-100-832). To generate embryoid-like bodies (EBs), iPSCs were harvested with Accutase and triturated into single-cell suspension. An Aggrewell-800 24-well plate (StemCell Technologies, #34811) was pre-washed with Anti-Adherence Rinsing Solution (StemCell Technologies, #07010) and DPBS, then 4 million cells were seeded per well in 2 ml of EB Medium: complete mTeSR + 0.05 ug/ml BMP-4 (PeproTech, #120-05) + 0.05 ug/ml VEGF (PeproTech, #100-20), + 0.02 mM SCF (PeproTech, #300-07) and centrifuged at 300 g for 3 min. EB Media was changed daily for three days by half medium change twice to avoid disrupting the formation of EBs. On day 4, EBs were collected in DPBS on a 37-um reversible strainer (StemCell Technologies, #27250). Harvested EBs were transferred to a tissue-culture treated 6-well plate (Falcon, #353224) in 3 ml of EB Differentiation media: X-VIVO 15 (Lonza Biosciences, #04-418Q) + 1% Glutamax (Life Technologies, #35050-061) + 1% Pen-strep + 0.1 ug/ml M-CSF (PeproTech, #300-25), + 0.025 ug/ml IL-3 (PeproTech, #200-03) + 0.055 M β-mercaptoethanol (Sigma, #M3148) at a target density of 70 EBs per well. A 2/3 EB Differentiation media change was done once to twice weekly depending on the growth rate. After several weeks, floating macrophage-like progenitors (pMacPres) emerging from cultures were isolated from the media through a 37-um cell filter and counted. 12.5 - 15K cells per 96-well, or 300K cells per 6-well tissue culture plates were seeded in Microglia Differentiation medium (Advanced DMEM-F12 (Gibco, #12634010) + 1x N2 supplement (Gibco, #17502048) + 1% Glutamax + 1% Pen-strep + 0.1 ug/ml IL-34 (PeproTech, #200-34; Biolegend Inc, #577904) + 10 ng/ml GM-CSF (PeproTech, #300-03) + 0.055 M β-merceptoethanol), to either 200 ul (96-well) or 3 ml volume (6-well) and grown at 37 C + 5% CO_2_ for 14 days.

### Phagocytosis assays

Phagocytosis assays were performed in 96-well plates (Corning, #3904) containing piMGs or iPSC-iMGs in a media volume of 100 ul. Human synaptosomes were thawed at room temperature and an equal volume of 0.1M sodium bicarbonate, pH 9 was added to the synaptosomes. pHrodo-Red (Invitrogen, #P36600, 6.67ug/ul) was added at a protein ug ratio of 1:2 (dye:synaptosome). The labeling reaction was incubated at room temperature for 1 hr in the dark. Synaptosomes were washed by adding 1 ml of PBS, pH 7.4 to the tube and pelleting at 12,000 rpm for 15 min. The wash solution was discarded and the labeled synaptosomes were resuspended in a volume of basal Advanced DMEM-F12 to a final concentration of 0.6 ug synaptosomes per ul. Labeled synaptosomes were sonicated in a Branson 1800 (Emerson, #M1800) at 40 kHz for 1 hr. During this time, some wells of microglia were treated with cytochalasin-D (Sigma, #C2618) to a final concentration of 1 μM to inhibit phagocytosis. Following sonication, the synaptosomes were added at a final concentration of 3μg/well. The assay was ended after 3 hrs by fixation with final 4% paraformaldehyde (Electron Microscopy Sciences, #15713S).

Additional assay details available in Supplemental Methods and Materials

## Results

### CYFIP1 is required for microglial-mediated synapse engulfment in a human in vitro model of synaptic pruning

Normal postnatal brain development results in surplus neuronal synapses that are either maintained or selectively eliminated, in a process peaking during early neurodevelopment as well as adolescence (16, 17), through microglial synaptic pruning.(18, 19) Feinberg first suggested that aberrant increased synaptic pruning during adolescence may contribute to the etiology of schizophrenia more than four decades ago.(20) While many neuronal genes that regulate the process of synaptic pruning have been implicated in schizophrenia, most notably the complement protein C4 (21, 22), little is understood about the role of schizophrenia-associated genes highly expressed in microglia (3, 23, 24), notably *CYFIP1* (10, 13, 25). We therefore utilized *in vitro* human models of microglia-mediated synaptic pruning, in conjunction with CRISPR-mediated editing to investigate possible roles for *CYFIP1* in this process.

We first tested the effect of *CYFIP1* knockdown using a human *in vitro* model of synaptic engulfment using cytokine-mediated reprogramming of human peripheral blood mononuclear cells (PBMCs) towards human induced microglia-like cells (piMGs) with IL-34 and GM-CSF.(22, 26) These piMGs show typical ramified morphology as well as express canonical microglial markers IBA1, P2RY12, CX3CR1 and TMEM119 (Figs. 1A-B). piMGs lacking *CYFIP1* expression were derived using CRISPR guide RNAs (gRNAs) directed to the *CYFIP1* gene resulting in frame-shift-induced truncation products of the CYFIP1 protein. Due to the primary nature of the PBMCs, unlike engineered cell lines that express Cas9 protein,(27-29) we adapted a modified protocol (30, 31) for CRISPR-mediated gene editing in PBMCs during piMG reprogramming using Cas9/sgRNA ribonucleoprotein (RNPs) complex delivery of purified recombinant Cas9 pre-complexed with either a synthesized single guide RNA (sgRNA) or a 2-piece CRISPR RNA:trans-activating RNA (cr:tracRNA) gRNA (Table S1) directed at *CYFIP1* resulting in efficient reduction of expression in the piMGs (Fig. 2A). The percent editing was determined by sequencing of the region, and the percentage of those resulting in frameshift edits (i.e., knock-out) predicted to result in functional truncation (Fig 2B). We observed >70% predicted knockout (frameshift inducing indels) of *CYFIP1* using both sgRNA and cr:tracRNA with no detectable alteration using the non-gene targeting negative control eGFP for each gRNA type (Fig. 2B).

**Figure 1.**
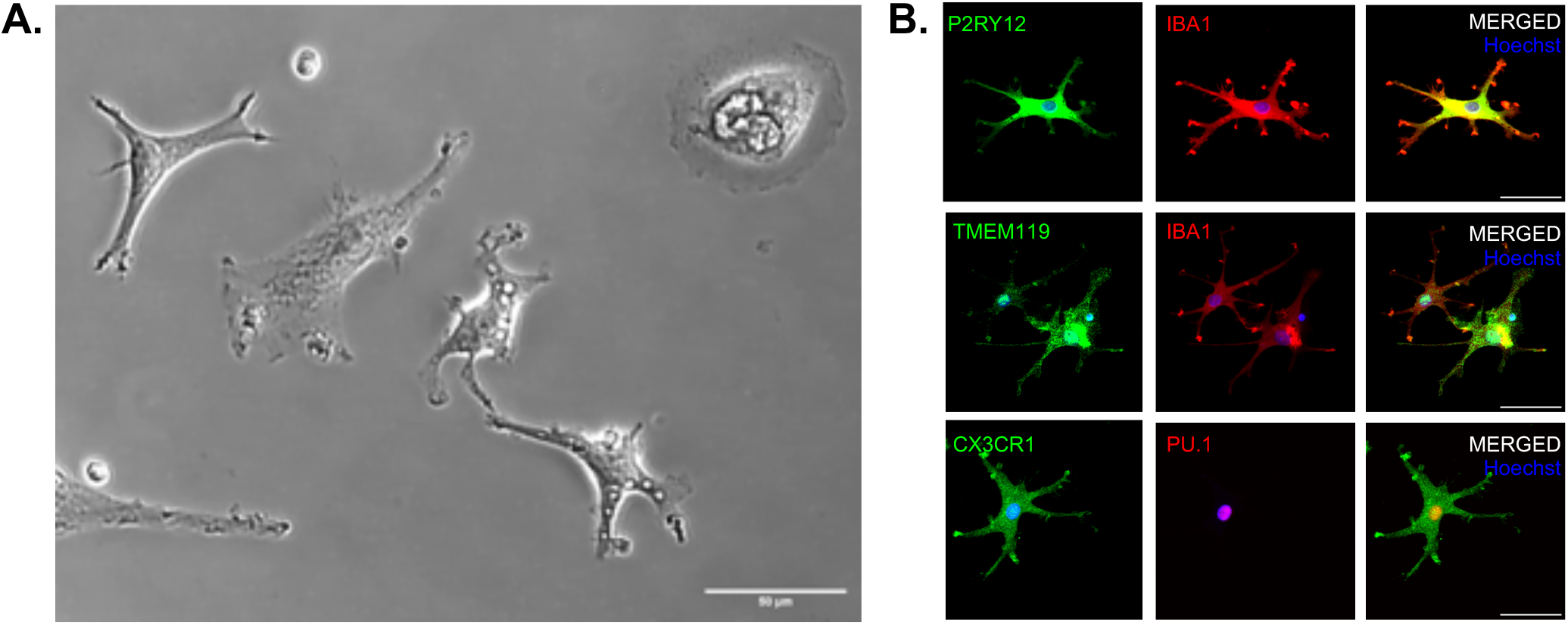
Characterizations of piMG cells. (a) Representative phase-contrast image of piMG cells captured during a live imaging session - Scale bar, 100 μm. (b) Confocal images of piMG cells stained for indicated canonical microglial markers P2RY12, IBA1, TMEM119, CX3CR1 and PU.1.

**Figure 2.**
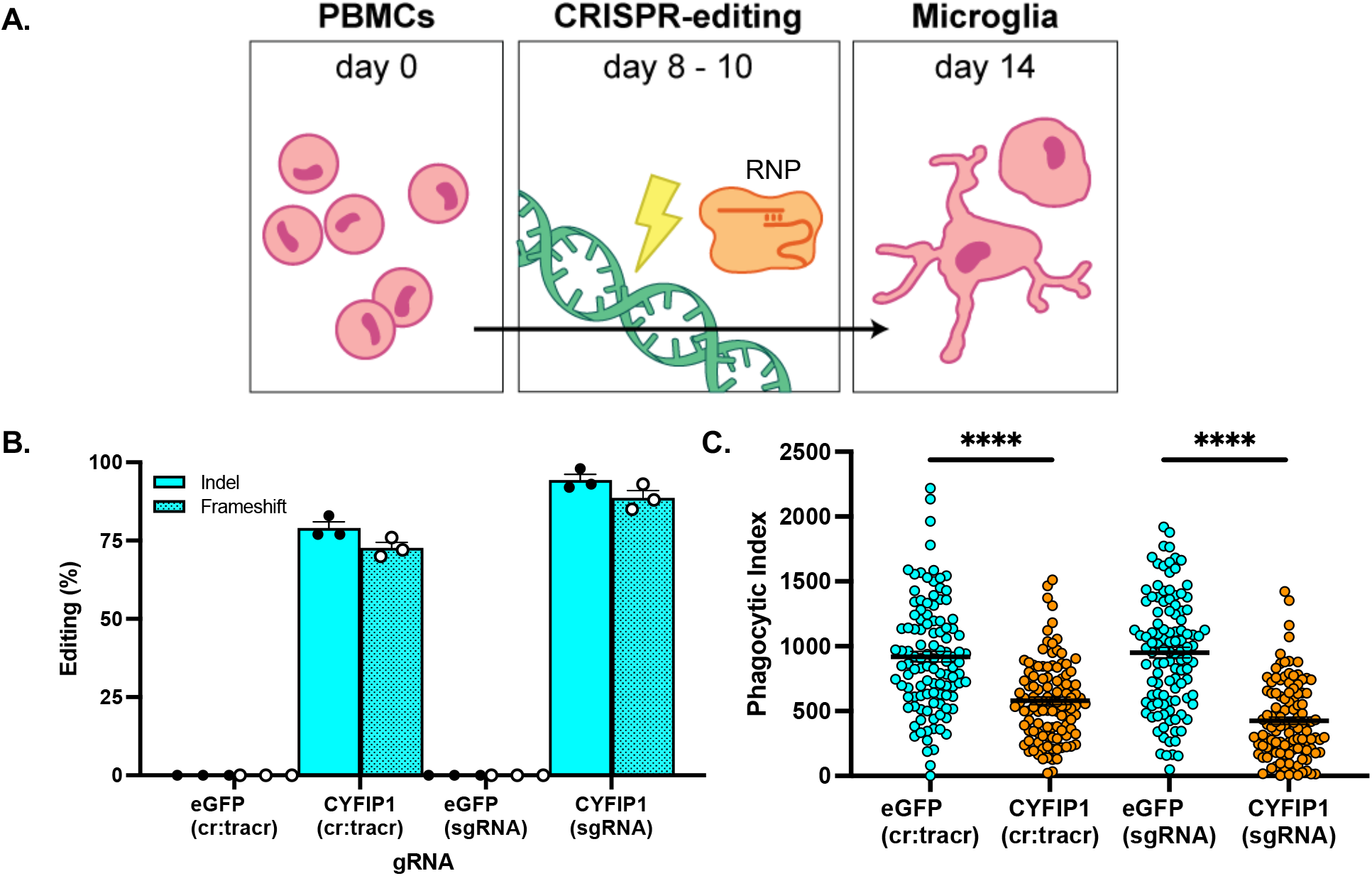
Gene editing of PBMCs to derive modified induced microglia. (a) PBMCs are harvested from large scale flask cultures and gene editing is performed during microglial reprogramming by nucleofection using pre-formed sgRNA/Cas9 ribonucleoproteins (RNPs) resulting in modified piMGs. (b) Editing efficiency with gRNA type as indicated for *CYFIP1* determined by sequencing (eGFP-RNP used as negative control). Percentages shown reflect editing events likely to yield functional KOs through frameshift. (c) Functional phagocytosis assays demonstrate correlative decrease in phagocytosis activity with *CYFIP1* knock-out efficiency. Each circle represents phagocytic index per image field. *n*=111 (37 fields in 3 wells) Line = mean, *** - *p* value<0.001

Functional phagocytosis assays were performed with piMGs nucleofected with *CYFIP1* or eGFP-directed cr:tracRNAs and sgRNAs, with human synaptosomes labeled with a pH-sensitive dye (pHrodo-red) that fluoresces upon specific engulfment into the acidic post-phagocytic phagolysosome compartment enabling robust quantification of synaptosome engulfment (phagocytic index) by confocal microscopy. *CYFIP1* deletion significantly decreased phagocytosis activity independent of gRNA type in the piMGs (Fig. 2C). These results support a direct role for *CYFIP1* function in microglia phagocytosis that, based on prior work (22, 26), can be extrapolated to mechanisms of human synaptic pruning.

### CYFIP1 expression is not essential for human iPSC pluripotency maintenance or capacity for microglia differentiation

While direct conversion of human blood monocytes towards microglia-like *in vitro* models has several advantages (32), including rapid and scalable generation of piMGs, direct conversion does not follow developmental ontogeny toward terminally differentiated endpoints. Microglia are of mesodermal origin, originating from the yolk sac during early development.(33-35) Protocols for deriving human iPSC induced microglia (iPSC-iMGs) that recapitulate this developmental ontogeny (15, 36-38), combined with manipulation of schizophrenia implicated genes such as *CYFIP1*, provides a means of characterizing additional roles in microglial viability, development, phenotype and function.

We further investigated the role of *CYFIP1* by creating clonal CRISPR-knockout iPSC-iMGs. iPSCs were transfected with CRISPR-RNPs followed by clonal selection verified by sequencing to ensure they are stably edited, allowing for subsequent clonal generation of iPSC-iMGs through a developmentally relevant process (15) to further identify potential effects on microglial genesis.

We used CRISPR-RNP nucleofection with cr:tracr gRNAs (Table S1) to knockout *CYPIF1*. Isogenic iPSC colonies were isolated after nucleofection (Fig 3A), expanded and sequenced at the *CYFIP1* locus to confirm editing in each clone (Fig. 3D). In total, we prioritized six clonal lines (Table S2) showing appropriate pluripotency marker analysis (OCT4, NANOG, TRA-1-60, Fig. S1) with confirmed normal karyotype (Fig. S2); two lines from an iPSC line derived from a healthy 29-year old male with homozygous frameshift indels resulting in predicted truncations through premature stop codon presentation (ko1-iPSC and ko2-iPSC, respectively); two lines that were not edited in the *CYFIP1* locus serving as isogenic controls (wt1-iPSC and wt2-iPSC) (Fig. 3D). In addition, to serve as a biological replicate, isogenic clonal *CYFIP1* knockout iPSC lines derived from a healthy 45-year old male were also chosen, one with a premature stop codon inducing homozygous frameshift indel (ko3-iPSC) and an isogenic control lacking editing (wt3-iPSC). *CYFIP1* gene expression was reduced by >100 fold in the edited lines (Fig. 3B) confirming knockout and likely transcript degradation. Deletion of CYFIP1 protein expression was confirmed by Western blot analysis (Fig. 3C). No noteworthy viability, doubling time or iPSC colony size (Fig. S3) differences were observed, demonstrating that *CYFIP1* expression is not essential for pluripotent maintenance or cellular viability.

**Figure 3.**
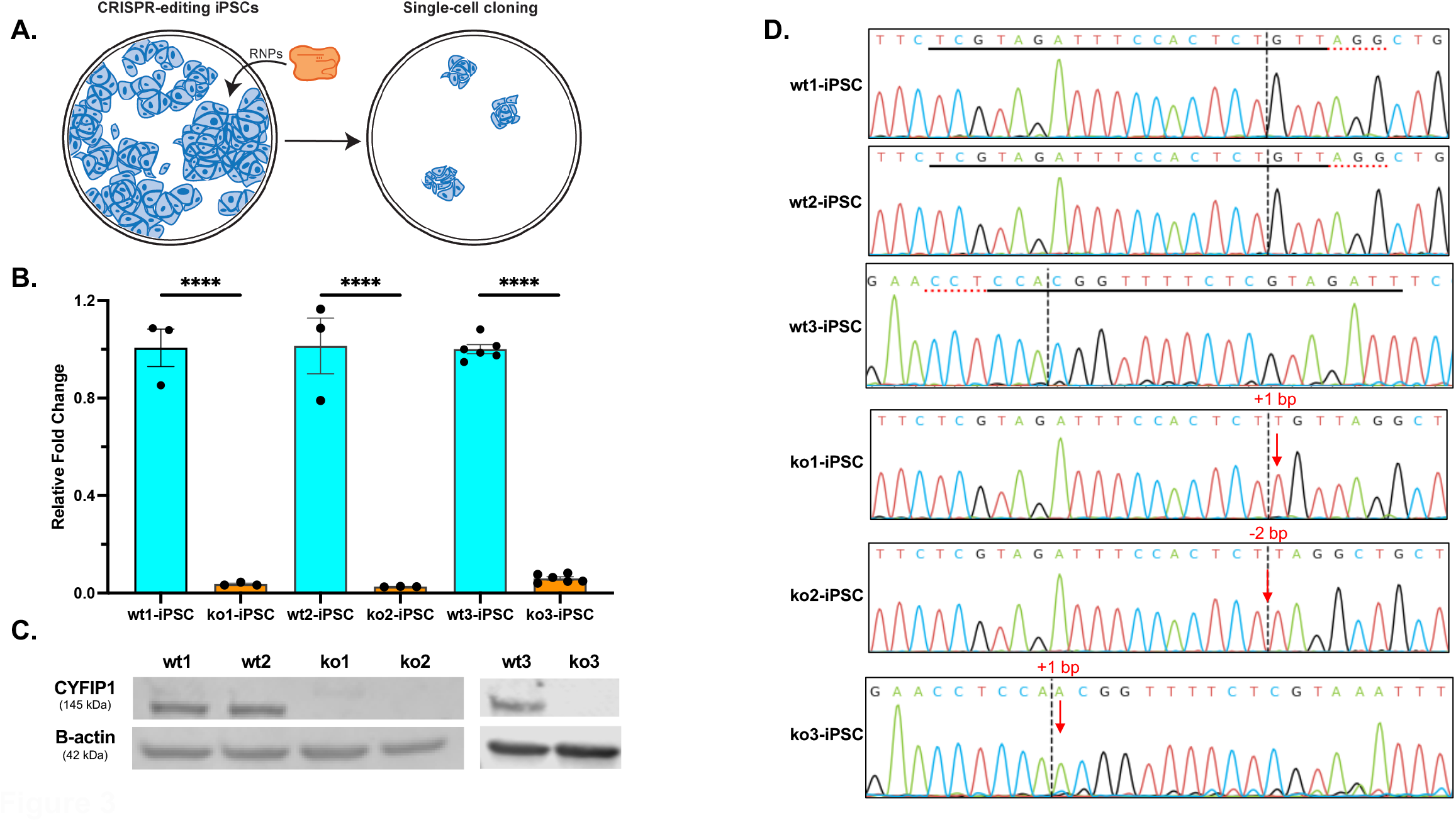
Derivation of CRISPR generated null human iPSC indicate *CYFIP1* not required for cellular viability (a) Generation of gene edited clonal iPSC lines using sgRNA/cas9 complexed ribonucleoprotein (RNP) nucleofection. (b) RT-qPCR quantification of *CYFIP1* expression normalized to housekeeping gene GADPH. Bar = mean, error = SEM of technical triplicates. *** - *p* value<0.001 (c) Confirmation of CYFIP1 protein deletion by Western blot analysis. (d) Clonal lines were expanded and editing (*CYFIP1*) was confirmed in each clone by sequencing.

Using modified procedures outlined in Haenseler *et. al*. to generate yolk-sac derived microglial precursors (pMacpres)(15, 39), we generated iPSC-iMGs from clonal *CYFIP1* KO iPSC lines (ko1-iMG, ko2-iMG, ko3-iMG) with unedited, isogenic iPSC control lines (wt1-iMG, wt2-iMG, wt3-iMG). Immunocytochemistry of conical microglial markers IBA1, P2RY12, CD11b and TMEM119 further confirms microglial identity in both wild type and *CYFIP1* KO iMG lines (Figs. 4A and B). Notably, *CYFIP1* KO did not significantly alter the expression of these markers (Fig. 4C). Loss of *CYFIP1* expression was confirmed in these differentiated iPSC-iMGs by RT-qPCR (Fig. 4D) and Western blot (Fig. 4E). Additionally, DNA sequencing demonstrated that the frameshift inducing indel in *CYFIP1* is retained upon iMG differentiation (Figs. S4 and S5) indicating a lack of reversion due to selective pressure during extensive proliferation and differentiation in culture.

**Figure 4.**
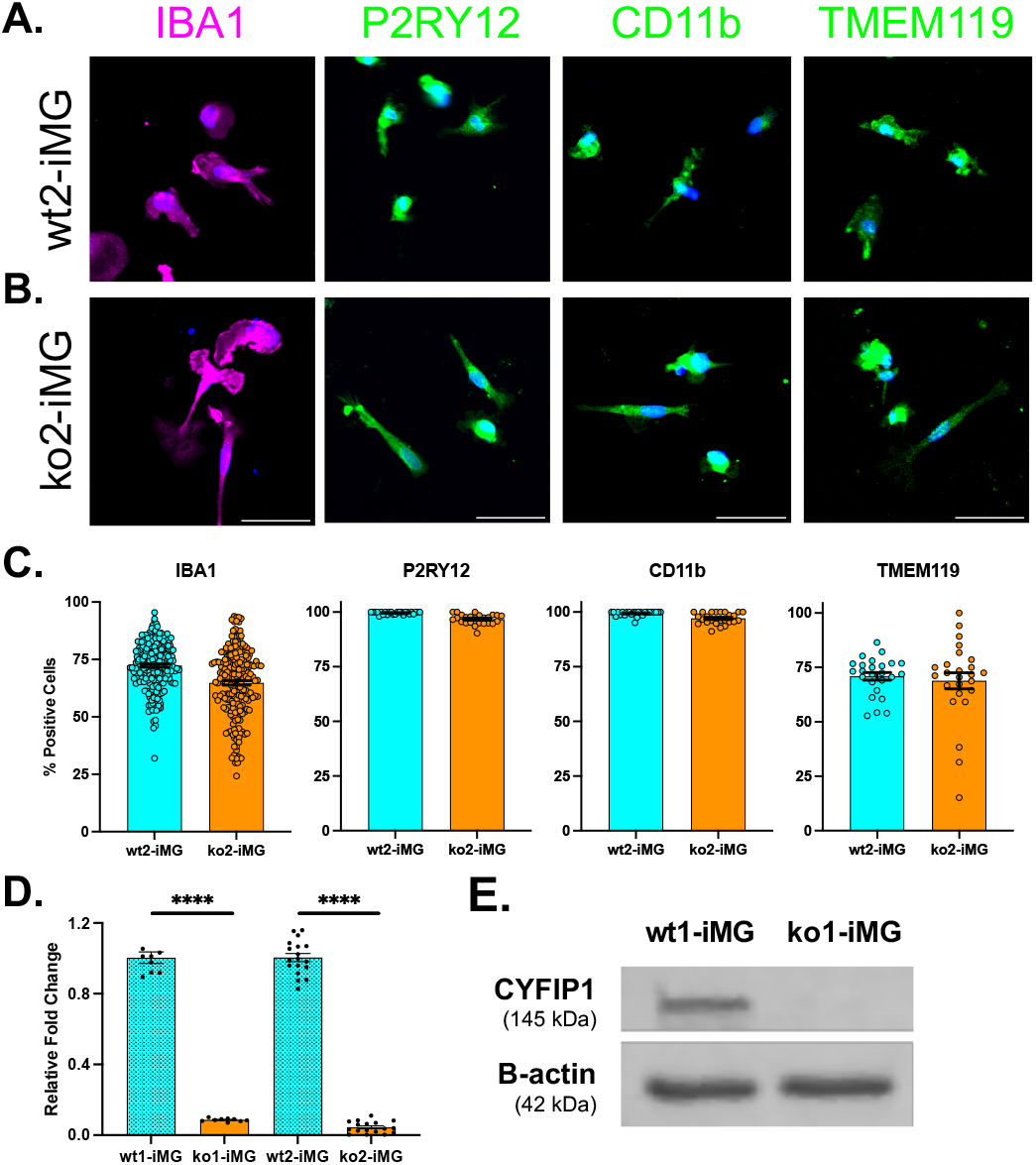
Characterization of iPSC-iMGs. Immunocytochemical analysis of microglial canonical markers (a) WT iPSC-hiMG (b) *CYFIP1* KO iPSC-MG images of iPSC-hiMG cells stained with nuclei (Hoechst) and indicated microglial markers IBA1, P2RY12, CD11b and TMEM119. Scale bar: 50 μm. (c) Quantitation of IBA1-positive immunostained cells as a percentage of total cells (nuclei), or as percentage of IBA1 positive cells for indicated microglial markers P2RY12, CD11B and TMEM119. Each circle represents mean of image fields (*n*=217 IBA1, *n*=25 for other markers). Bars indicate mean, error bars indicating SEM. (d) Quantitation of *CYFIP1* gene expression in indicated WT and KO lines after iMG reprogramming, error bars indicating SEM (n=9 replicates for wt1/ko1; 18 replicates for wt2/ko2), **** - *p* value<0.0001 (e) Western blot analysis of CYFIP1 protein expression in indicated WT and KO iMG lines.

While embryonic failure demonstrated by murine homozygous *CYFIP1* knockout suggests essential global roles in the early stages of embryogenesis (6, 14), our demonstration that human iPSCs lacking *CYFIP1* expression retained pluripotency and the capacity to form intermediate microglial precursors and functional microglia *in vitro* suggests that the expression is not required in this terminal differentiation developmental process.

### Requirement of microglial CYFIP1 expression for engulfment of synaptic structures in iPSC-derived microglia supports a potential role in microglial-mediated synaptic pruning

While our data supports the premise that deletion of *CYFIP1* is not required for terminal microglial differentiation from human iPSCs, we sought to further confirm our observation of reduced synaptosome engulfment in clonal *CYFIP1*-null iPSC-iMGs using our *in vitro* synaptic pruning assay. iPSC-derived microglia precursors (pMacpres)(15) from wild type and clonal *CYFIP1* null lines were seeded into 96-well plates in final maturation media containing IL-34 and GM-CSF for 14 days. Phagocytosis assays were initiated with the addition of excess pHrodo-red labeled synaptosomes as described above for PBMC-iMGs. iPSC-iMGs were incubated with synaptosomes for 3 hours, after which cells were fixed and stained for subsequent quantitation of the phagocytic index of synaptosome engulfment. While all clonal lines exhibited synaptosome uptake indicative of phagocytic function, the *CYFIP1*-null iMGs demonstrated significantly reduced engulfment as compared to isogenic controls (Fig. 5). This reduction was observed across lines generated from two different donor sources (29 and 45-year old healthy male donors) and gRNAs (Tables S1 and S2) demonstrating a robust phenotype of decreased synaptosome engulfment upon *CYFIP1* ablation, consistent in multiple donor lines and *CYFIP1* frame shift functional deletions.

**Figure 5.**
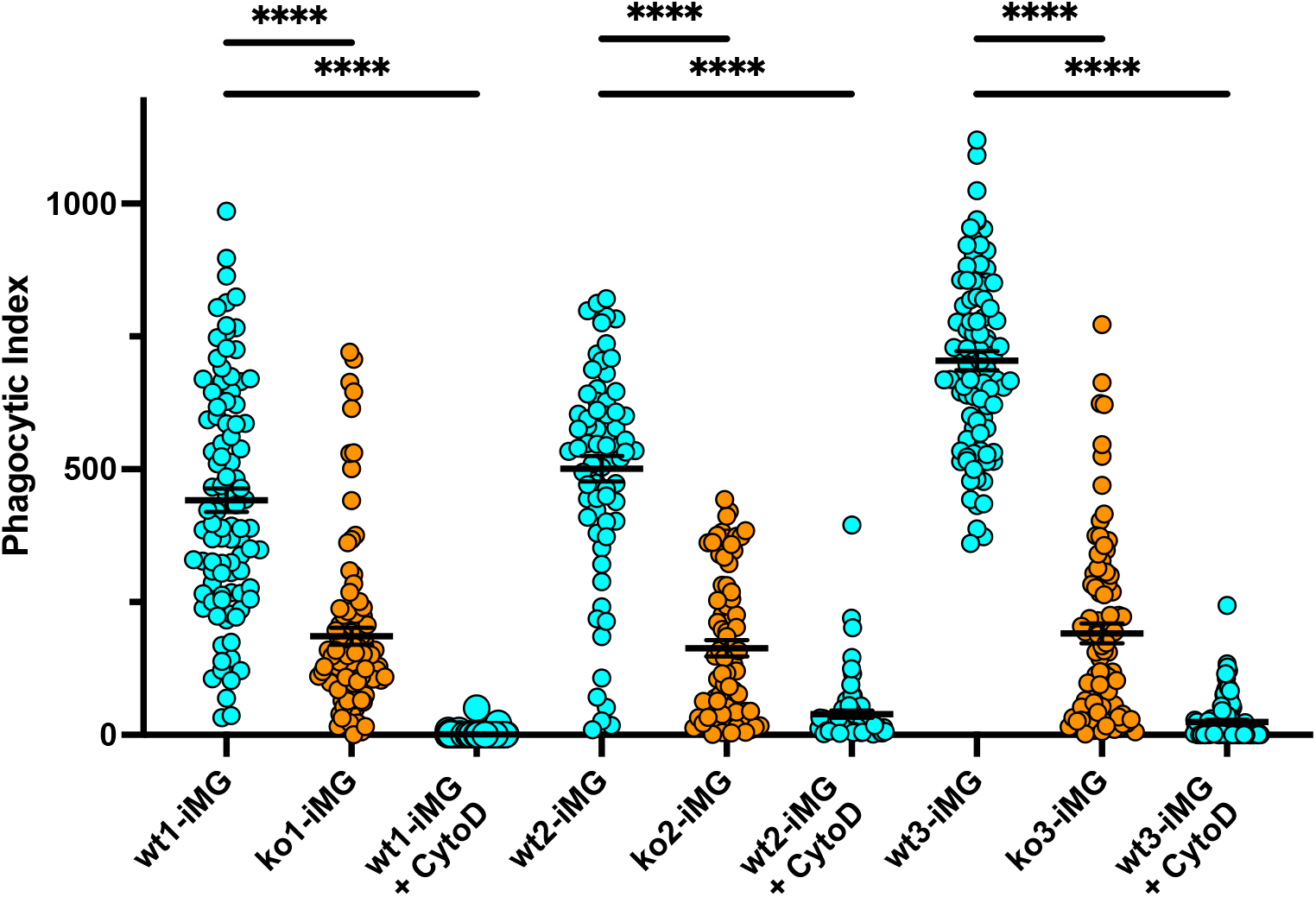
Reduced phagocytosis of synaptic structures in iPSC-hiMGs confirms potential role of *CYFIP1* in synaptic pruning. Functional synaptic engulfment characterization of *CYFIP1* WT and knockout hiMG lines by *in vitro* synaptosome engulfment. Each circle represents phagocytotic index per image field across 3 technical replicate wells = area of red engulfed synaptosomes/cell count per field. cytoD - pos. control inhibitor. Bar = mean, error = SEM, **** - p value<0.0001.

In agreement with the demonstration of reduced synaptic engulfment upon *CYFIP1* silencing in our orthogonal piMG model, these observations suggest that *CYFIP1* does not have an essential role in microglial differentiation, though it has essential direct or indirect functional roles in synaptosomal engulfment in terminally differentiated microglia, further supporting a role in functional synaptic pruning.

### Loss of CYFIP1 in human iPSC-derived microglia results in altered morphology and motility

*CYFIP1* is highly expressed in microglia in the human CNS,(10-12) and the protein has been shown to mediate interactions between the WAVE regulatory complex (WRC) and Arp2/3 complexes (40), allowing the Arp2/3 complex to nucleate the branching and extension of actin filaments (41, 42) which in part regulates actin cytoskeleton remodeling during cellular processes involved in morphology and migration that have been implicated in the regulation of murine microglial state.(43) We sought to further characterize potential functional effects by phenotypic comparison of wild type and *CYFIP1* loss-of-function clonal iPSC-iMGs including morphology (e.g., resting-state arborization) and motility (e.g., surveillance) that may be related to the activation state,(44) recognizing that microglial activation is not a unidimensional process. That is, we examined phenotypes in *CYFIP1* KO iPSC-iMG lines to detect effects on morphology and motility.

Morphological analysis was performed using an automated image analysis pipeline in CellProfiler (45) to determine morphometric measures of solidity and eccentricity for IBA1-stained iPSC-iMGs (Fig. 6). We have shown that these parameters are sufficient to categorize individual cells (Fig. 6A) into ramified (low solidity, high eccentricity), amoeboid (high solidity, low eccentricity) and bipolar, or rod shaped (mid solidity, mid eccentricity) morphotypes in a high-throughput fashion. Using this procedure, we measured these parameters for wild type and *CYFIP1*-null iPSC-iMGs lines wt1-iMG, ko1-iMG, wt2-iMG and wt2-iMG. (Fig 6) From these analyses, we observed that loss of *CYFIP1* resulted in an overall morphology shifted toward a more ramified phenotype compared to their isogenic controls as indicated by significantly (*p < 0*.*0001*) lower solidity (Fig. 6B) and higher eccentricity (Fig. 6C) morphometric values.

**Figure 6.**
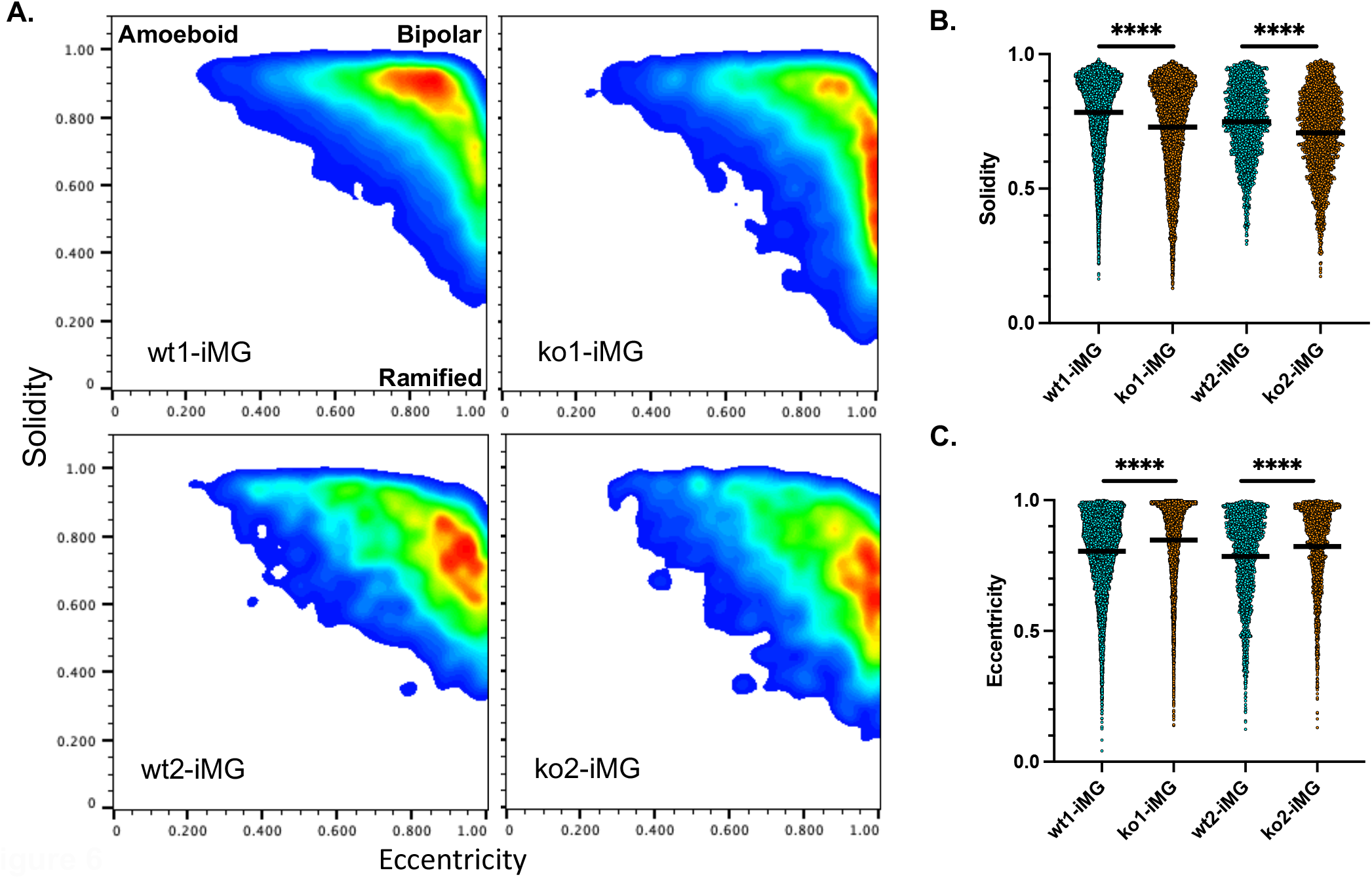
Loss of *CYFIP1* in iPSC-iMGs results in altered morphology. (a) Morphology smoothed density plots (solidity vs. eccentricity). WT cells exhibit a less ramified morphology than *CYFIP1*-KO, suggestive of a less activated phenotype. Graphical representations of (b) solidity and (c) eccentricity. Each circle – individual cell. Bar indicates mean +/- SEM, **** - *p* value <0.0001. Number of cells *n*=6502 (wt1-iMG), 10601 (ko1-iMG), 2204 (wt2-iMG), 2440 (ko2-iMG).

*CYFIP1* has been shown to be involved in cellular motility via its control of actin dynamics (40-42), thus we hypothesized that it may be required for proper motility required during microglial migration during early CNS development as well as surveillance in the postnatal brain.(46) We measured cellular motility using real-time imaging for 5 hours (movies S1 and S2). Loss of *CYFIP1* function resulted in a striking significant reduction in cellular motility as demonstrated comparing lines wt2-iMG, ko2-iMG, wt3-iMG and ko3-iMG (Fig. 7A). Both the mean distance traveled during real-time imaging (Fig. 7B), as well as mean velocity (Fig. 7C), were shown to be significantly (*p*<0.0001) reduced in the *CYFIP1* KO iMGs compared to their isogenic controls.

**Figure 7.**
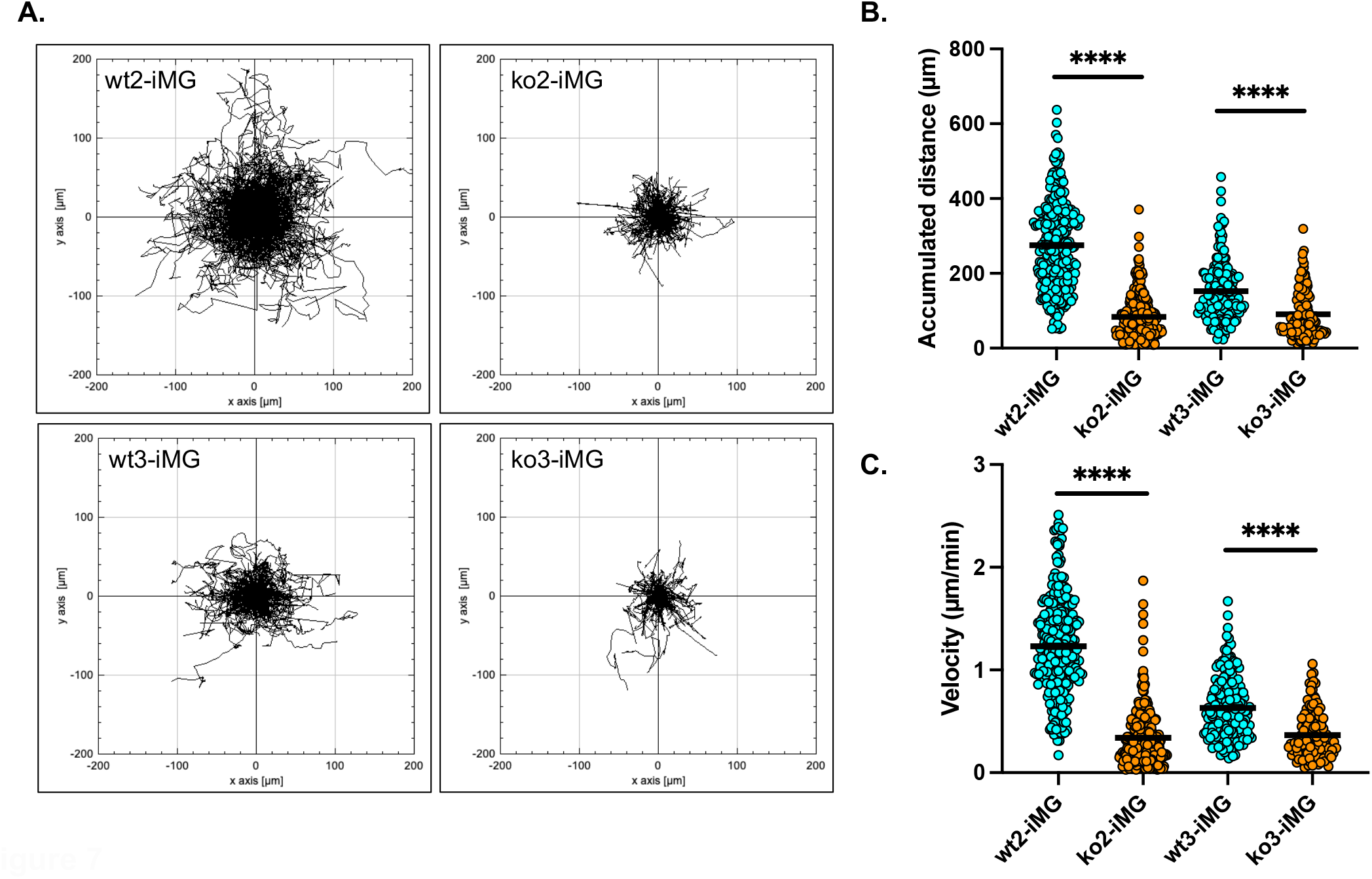
Loss of *CYFIP1* in iPSC-iMGs results in reduced motility. (a) Migration plots by real-time imaging every 5min for 5 hours with each track from an individual cell aligned at starting point. Axis units in mm from starting point. (b) Accumulated distance (um) and (c) migration velocity (um/min) over 5 hours are greatly reduced in *CYFIP1*-KO iMGs. Bar indicated mean. **** *p* value <0.0001. Numbers of cells *n*=273 (wt2-iMG), 288 (ko2-iMG), 190 (wt3-iMG), 128 (ko3-iMG).

## Discussion

Most efforts to understand the pathophysiology of neurodevelopmental and neuropsychiatric disorders have focused on studies of aberrant neuronal function, overlooking the central role of microglia in normal neurodevelopment. Microglia populate the brain very early in embryonic development (47-49) and are critical in coordinating neuronal progenitor populations (50, 51), neurogenesis (52-54) and the organization of synaptic networks through the selective process of synaptic pruning. (18, 55, 56)

To address this gap, we investigated the role of the *CYFIP1* gene, which resides within the 15q11.2 region that has been directly implicated in the etiology of neurodevelopmental disorders. (3, 23, 24) While its role in neuronal function has been extensively characterized, *CYFIP1* expression in the CNS is actually highest in microglia (10-13), where its role is not understood. Specifically, we focused on the effects of *CYFIP1* in human models of synaptic pruning, in light of the microglia-mediated role of pruning in schizophrenia (16, 20, 57, 58) as well as increased synaptic density (hypo-pruning) in autism and epilepsy.(59) Prior studies have demonstrated that microglia play a key role in the selective pruning of synapses and maintenance of proper brain connectivity during adulthood (60-63) and aberrant microglial function has been implicated in several neurodevelopmental and neuropsychiatric disorders.(32, 59, 64)

We found that loss-of-function of *CYFIP1* though CRISPR engineering in orthogonal *in vitro* studies using both human PBMC-derived microglia (piMGs) and iPSC-derived microglial cellular models (iMGs) resulted in robust reduction of phagocytosis of human synaptosomes in *in vitro* models of synaptic pruning.(22, 26, 32). Together, these models suggest a critical role for microglial *CYFIP1* function that can be altered through aberrant genetic contributions that effect normal expression and/or functional levels though haploinsufficiency or regulatory variants.(1, 65, 66)

In addition to their central role in synaptic pruning, microglia exhibit dynamic morphological plasticity during normal development(67) as well as in response to stimuli such as injury and disease.(68, 69) Microglia morphology varies significantly across different brain regions, though during homeostatic conditions microglia typically display a complex ramified morphology indicative of a migratory surveillant state, making transitory contacts with neurons, synapses, blood vessels, and astrocytes.(46) Morphological shifts towards a rounded amoeboid morphotype has been associated with activation due to external stimuli such as brain injury (69), proinflammatory stimulus and disease states, while rod-like bipolar microglia have been demonstrated to be highly associated with neurons and implicated in synaptic stripping.(70, 71) These morphotypes are also highly influenced by developmental stage: embryonic and early post-natal microglia tend to exhibit a more amoeboid form, whereas they can adopt a highly dystrophic form as a result of extended age or as a result of neurodegenerative disease.(72) Thus, various microglial morphotypes in these different contexts suggest a form-function relationship in which morphology and expression of various markers can be used as a proxy for functional state.(73, 74) CYFIP1 protein has been demonstrated to directly interact with the Wiskott-Aldrich syndrome protein member 1 (WAVE1) regulating dendritic cytoskeletal modulation of ARP2/3-mediated actin branching.(40-42) Our morphological characterization in *CYFIP1* loss-of-function iPSC-derived microglia implicates a direct role in the cytoskeletal processes involved in proper function. In agreement with the observed reduction in phagocytic function, we observed that the loss of *CYFIP1* expression in these iMGs results in increased ramification consistent with a less activated state.(67, 73, 74)

Microglial motility into and within the CNS is also essential to proper neurodevelopment and maintenance functions.(75) These cells must first properly migrate from the primitive yolk sac to populate the developing brain during early development to become permanently brain-resident throughout life.(49) Population of the developing CNS by microglial precursors occurs during defined developmentally timed critical phases regulated by both migration of peripheral sources and proliferation.(49, 75) In addition to populating the developing brain during this early migration and parallel proliferation, proper timing and brain region specificity is essential. Additionally, motility of the microglia in postnatal and adult brain are required for the function of surveillance, in particular during periods of synaptic development and refinement.(59, 76) Our demonstration that *CYFIP1* is essential for microglial motility, likely through its essential role in cytoskeletal actin polymerization, further suggests that influences on its expression have functional impacts for critical microglial functions in developmental migration and adult surveillance.(46, 49, 75)

Furthermore, we demonstrated that *CYFIP1* is not required for the maintenance of pluripotency during expansion of human iPSCs or the ability to be differentiated toward microglia. Prior work indicated that *CYFIP1* loss is lethal in murine embryos, making its role in early development challenging to determine. Murine studies had demonstrated the ability to form early embryonic homozygous *CYFIP1* knockout murine blastocysts, but resulted in developmental failure suggesting the critical requirement for *CYFIP1* in later developmental post-blastula roles (6, 14). Our results suggest that microglial loss of *CYFIP1* is unlikely to mediate such failure.

Taken together, our results showed multiple effects of *CYFIP1* loss-of-function on microglial phenotypes beyond diminished phagocytosis, including increased ramified morphology suggestive of a less activated, resting state and greatly reduced motility. While microglia differentiation is grossly preserved, alterations in expression level (e.g. through 15q11.2 (BP1-BP2) CNV haploinsufficiency) or functional variants likely impact proper functioning of microglia. These functional impacts can contribute during early developmental population of the brain, synaptic pruning during adolescence or normal homeostatic surveillance and maintenance contributing to neurological disease etiology such as schizophrenia resulting from microglia dysfunction. Our results do not detract from the importance of understanding the effects of neurodevelopmental risk genes on neuronal function, but underscore the need to further investigate these genes in microglia as well. They support the potential utility of investigating modulators of microglia as potential strategies to modify disease course in a range of neurodevelopmental disorders.

## Acknowledgements

This work was supported by R01MH120227 (Dr. Perlis)

## Figure Legends

**Movie S1**. Loss of *CYFIP1* in human iPSC-derived microglia results in reduced motility. Representative field of real time imaging of motility taken every 5 min per frame over 5 hours for wild type line wt2-iMG.

**Movie S2**. Loss of *CYFIP1* in human iPSC-derived microglia results in reduced motility. Representative field of real time imaging of motility taken every 5 min per frame over 5 hours for *CYFIP1* KO line ko2-iMG

## Supplemental Methods and Materials

### iPSC-derived neural differentiation for synaptosome isolation

iPSCs were reprogrammed from fibroblasts and used to derive neural progenitor cells, which were differentiated into neural cultures, as previously described.(22, 77) Neural progenitor induction from iPSCs (healthy control male donor line PSC-01-179) was performed with iPSCs cultured in E8 medium (Gibco) on Geltrex coated six-well plates and passaged using 50 mM EDTA and trituration with ROCK inhibitor (10 mM Thiazovivin; Stemgent). Neural progenitor cells (NPCs) were derived from these iPSCs using Neurobasal Medium (Thermo Fisher Scientific, #21103049) with 1X Neural Induction Supplement (Thermo Fisher Scientific, #A1647801), expanded using a neural expansion medium, and purified by double sorting using MACS against CD271 and CD133. NPCs were validated by immunocytochemistry for canonical NPC markers including Nestin, SOX1, SOX2, and Pax-6. Neural differentiation was initiated on Geltrex-coated T1000 5-layer cell culture flasks (Millipore Sigma, #PFHYS1008) and grown in neuronal differentiation medium (Neurobasal media (Gibco, #21103049) supplemented with 1X each (N2 supplement (Stemcell Technologies SCT, #7156), B27 supplement without Vitamin A (Gibco, #12587010), Non-essential amino acids (NEAA Gibco, #11140050), penn/strep), 1 uM Ascorbic Acid, 10ng/mL BDNF and GDNF (Peprotech) and 1ug/ml mouse laminin (Sigma, #L2020) for 8 weeks.

### Large scale generation of synaptosomes from iPSC-derived neural cultures for use in models of synaptic pruning screening assays

Neural cultures were differentiated from iPSC-derived neuronal cultures in high-yield T1000 multilayer flasks as we have previously described.(78) Synaptosome isolation from harvested cultures was performed by sucrose gradient was adapted for iPSC-derived differentiated neural cultures from previously described protocols.(79-81) Cells were harvested in 1X gradient buffer (ice-cold 0.32M sucrose, 600mg/L Tris, 1 mM NaH_3_CO_3,_ 1mM EDTA, pH 7.4 with added HALT protease inhibitor – ThermoFisher, #78442), homogenized using a dounce homogenizer and centrifuged at 700 *g* for 10 minutes at 4°C. The pellet was resuspended in 10ml of 1X gradient buffer (transferred to a 30 mL and centrifuged at 15,000 *g* for 15 minutes at 4°C. The second pellet was resuspended in 1X gradient buffer and slowly added on top of a sucrose gradient with 1X gradient buffer containing 1.2M (bottom), 0.85M (middle) sucrose layers. The gradient and cell mixture was centrifuged at 26,500 rpm (∼80,000 *g*) for 2 hours at 4°C, with the brake set to “slow” so as not to disrupt the final bands. The synaptosome band (in between 0.85M and 1.2M sucrose) was removed with a 5-mL syringe and 19Gx1 ½” needle and centrifuged at 20,000 *g* for 20 minutes at 4°C. The final pellet was resuspended in an appropriate volume of 1X gradient buffer with 1mg/mL bovine serum albumin (BSA) with protease and phosphatase inhibitors, aliquoted and slowly frozen at -80°C. Protein concentration was measured by BCA.

### Sanger sequencing

To screen for clonal iPSCs, cell lysates were generated by adding lysis buffer, 20 mM Tris pH 8, 0.1% Triton X-100 (Sigma, cat#T8787), and 60 ng/ul Proteinase K (NEB, #P8107S), directly onto live cells and scraping to collect. Lysate solutions were then heated to 65 C for 10 min, then 98 C for 2 min, and stored at -20 C if needed. PCR reactions (50 ul) were performed using the Phusion High-Fidelity PCR Kit (NEB, E0553L) following manufacturer’s protocol using 2 - 5 ul of lysate, and primers specific to the genomic loci of interest (Table S1). Reactions were run as follows: 98 C for 10 s, 35 cycles of [98 C for 5 s, 63 C for 10 s, 72 C for 30 s], and 72 C for 3 min for final extension. PCR amplicons were purified using the QIAquick PCR Purification Kit (Qiagen, #28106) and quantified with a Qubit3 fluorometer (Invitrogen, #Q33216) using the High Sensitivity dsDNA Quantitation kit (Invitrogen, #Q32854).

For validation of iPSC, microglia, astrocyte or neuronal lines, cells were harvested from culture by removing the media, washing with DPBS (Life Technologies, cat#14190), adding 1 ml of Accutase solution, and incubating at 37C for several minutes until the cells unadhered. Cells were harvested into a conical and pelleted at 300 g for 5 min. The media was discarded, and cell pellets were immediately processed with the QIAamp DNA Mini kit (Qiagen, #51306). Purified DNA was quantified with a Qubit3 fluorimeter using the High Sensitivity dsDNA Quantitation kit. 2 – 5 ng of purified gDNA was used as input for PCR under the following cycling conditions: 98 C for 10 s, 32 cycles of [98 C for 5 s, 63 C for 10 s, 72 C for 30 s], and 72 C for 3 min for final extension. Amplicon purification was performed as previous described then sent to Genewiz for sanger sequencing. Gene editing analysis of sanger traces was done using Synthego’s ICE webtool (https://ice.synthego.com/).

### Western blot

Clonal iPSC cell stocks (∼1 million/vial) were quick thawed at 37 C then immediately transferred into an Eppendorf tube and pelleted at 300 g for 5 min. The freezing solution was discarded, and the pellet was lysed using 30 ul of 1x RIPA buffer (Boston Bioproducts, # BP-116T) + PhosSTOP EASY phosphatase inhibitor (Roche, #04906837001), + cOmplete protease inhibitors (Roche, #04693159001). The lysis reactions proceeded for 15 min on ice, then were pelleted at 15,000 rpm at 4 C. The protein solution was carefully removed without disrupting the pellet and moved into a new Eppendorf tube. Total protein concentrations were calculated using the Pierce BCA Protein assay (Thermo Scientific, #23225) following manufacturer’s protocols.

A Bio-Rad Electrophoresis Cell (#1658004) was used to run the mini-PROTEAN gel (BioRadand #456-1096) in 1x Tris-Glycine-SDS running buffer (Boston BioProducts, #BP-150). Protein samples were prepared by mixing 25 ug total protein with 6x Loading Dye (Boston BioProducts, # BP111R) diluted to 1x in DI water, and heated to 95 C for 2 min immediately prior to loading. 4 ul of Precision Plus Protein Dual Color Standards (Bio-Rad, #1610374) were also run to visualize protein migration. The samples were run at 60 V for 15 min, then 100 V for 1.5 hrs.

Transfer steps utilized the BioRad Mini PROTEAN Tetra Cell system. 10x Transfer buffer (Boston BioProducts, #BP-190), was diluted to 1x final with DI water and 20% methanol. An Immun-Blot PVDF membrane (Bio-Rad, #1620177) was activated in 100% methanol for 15 s, submerged in miliQ water for 2 min, then finally in Transfer buffer for 5 min. All materials used in the transfer were likewise submerged in Transfer buffer including the Extra Thick Blot Filter paper (Bio-Rad, #1703967), sponges, and sandwich cassette. Following assembly, an ice pack was placed into the reservoir and the transfer occurred at 125 V for 1 hr. Membranes were blocked using LICOR Intercept Blocking Buffer (LI-COR, #927-70001) and rocked for 1 hr at 4 C. Blocking buffer was replaced with primary antibody solutions diluted 1:1000 in Blocking Buffer + 0.2% Tween-20 (Sigma, #P2287) (Table S1) Rabbit anti-CYFIP1, Abcam, #ab156016; mouse anti-beta-Actin, Abcam, #ab8226) and rocked at room temperature for 1 hr. Membranes were washed with PBS + 0.01% Tween-20 by rocking for 3 min at room temperature for a total of three washes. Secondary antibodies IRDye 800CW Donkey anti-Rabbit IgG, LI-COR, #926-32213; IRDye 680RD Donkey anti-Mouse IgG, LI-COR, #926-68072) were prepared at a 1:15,000 dilution in 50% blocking buffer + 50% [DPBS + 0.01% Tween-20] + 0.02% SDS and added to the membrane and rocked at room temperature for 45 min. Additional final washes were done as previously described before imaging with the LICOR Odyssey Clx.

### RT-qPCR measurement of gene expression

RNA was extracted from live cells by first washing the cells with media, aspirating the wash, then adding on Lysis buffer using Qiagen, miRNeasy kit for several minutes, followed by scraping. Subsequent RNA purification was performed following manufacturer’s protocols and eluted in recommended volume. RNA was quantified using the Qubit3 system with the High Sensitivity RNA kit (Invitrogen, # Q32851). 200 ng of RNA was treated with 1 ul of dsDNAse in the provided dsDNAse buffer (Thermo Scientific, #EN0771) and incubated at 37 C for 10 min, then 1 ul of 0.1M DTT was added and reactions were incubated at 55 C for 5 min, put on ice and quantified again using the Qubit3. 50 ng of RNA were used as input for reverse transcription using the SuperScript™ III First-Strand Synthesis System (Invitrogen, #18080051) as per two-step protocol with RNAseH. Control reactions included complete reaction conditions with either no RNA template or no reverse transcriptase. RT-qPCR reactions were performed in triplicate for both the gene of interest and loading control, which were multiplexed. Each 10 ul qPCR reaction occurred in a 384-well plate and contained 5 ul of 2x TaqMan™ Fast Advanced Master Mix (Invitrogen, #4444557) + 3 ul of diluted cDNA (∼ 2 ng) + 0.5 ul of each 20x Taqman kit (Table S1) in water. Reactions were cycled in a Roche LightCycler 480 II as follows: 50 C for 2 min, 95 C for 2 min, 40x cycles of [95 C for 1 s, 60 C for 20 s + single acquisition], 40 C for 10 s. Data was exported and analyzed such that values Ct values of 35 and higher were considered noise inherent in this assay. Each gene of interest Ct value was normalized to the loading control in the same well, then to the average of the triplicate values from the experimental control. Fold change was calculated as 2^(-(deltadeltaCt)).

### Immunofluorescence

Live cells were fixed with 4% paraformaldehyde for 15 min at room temperature. Cells in a 96-well format were washed with 100 ul of Wash Buffer, PBS + 0.5% FBS, three times. Then 100 ul of Block and Permeabilization Buffer, PBS + 0.5% FBS + 0.3% Triton-X, were added per well. Wells were incubated at room temperature for 1 hr, then washed three times with Wash Buffer. Primary antibodies (Table S1) were diluted in Antibody buffer, PBS + 0.5% FBS + 0.1% Triton-X, added to the wells and incubated for 1 hr at room temperature or overnight at 4 C. Wells were washed three times with Wash Buffer. Secondary antibodies (Table S1) were diluted in Antibody Buffer and added to wells and incubated for 45 min at 4 C. After a final three washes, the wells were imaged with an IN Cell Analyzer 6000 automated confocal microscope (Cytiva) at 20.

### High Content Image Analysis

CellProfiler (Version 4.2.4) (45) was used to identify iMGs for the purpose of quantifying percent marker positive cells, cell morphology, and phagocytosis of synaptosomes. The key modules of the pipeline used: Images from blue, green, red, and far-red channels were matched by field in NamesAndTypes. First, the images were background corrected using CorrectIlluminationCalculate and CorrectIlluminationApply. iMGs were identified from IBA1 staining in the far-red channel using IdentifyPrimaryObjects, with nuclei and other markers using the same module applied to the blue and green channels, respectively. iMGs were then filtered by expected area range using FilterObjects. The pHrodo red labeled synaptosome images were masked to the cell and nuclei objects so that only red signal within cells was included, with any possible signal outside the cells or within the nuclei omitted. Using those masked images, synaptosomes were identified using the IdentifyPrimaryObjects module. This module specifically uses the adaptive Otsu thresholding strategy, which increased accuracy. Further, we ensured background noise would not be included by running the pipeline with images from wells without the addition of synaptosomes and making sure there was no segmentation of red signal. At this point all necessary objects are identified and measurement modules follow, including MeasureImageAreaOccupied, MeasureObjectSizeShape, and MeasureObjectIntensity. Finally, RelateObjects was used to organize data into the proper relational formatting, and data was exported using ExportToSpreadsheet.

We next used RStudio to analyze the exported metadata from CellProfiler. In short, cell and synaptosome metrics were compiled by image field to quantify phagocytic index, which is defined as the area of pHrodo red signal divided by number of cells per image. Marker percentage was also done at the image field level, where percent IBA1 positive cells was found by dividing the number of these object by the total nuclei count. Further marker percentage was found by dividing number of cells with IBA1 that also contain a second marker by the number of cells positive only for IBA1. For morphology analysis, individual cell morphology, including solidity and eccentricity data, were compiled and compared between cell line groups.

### Real time cell-imaging and data analysis

Live-imaging of microglia cultures was performed in the Sartious Incucyte (Zoom). For assays of motility, microglia were imaged at 20x every 5 mins for 5 hrs. Image sets were exported from the software as 8-bit .tif files and imported into FIJI image analysis software (82) version 2.3.0/1.53q as an Image sequence. Using the stabilization plug-in for ImageJ, Image Stabilizer (83), the sequences were stabilized to offset changes in registration inherent in live-imaging and saved as a new image sequence.

### Motility assays

Real time label-free phase contrast imaging was performed on the Sartious Incucyte (Zoom) at 20x magnification imaging 4 fields (wt2-iMG and ko2-iMG) or 3 fields (wt3-iMGs and ko3-iMG) every 5 min for 5 hours. Images were stabilized using Image Stabilizer.(83) To track cells in the sequence, stacked images were combined to a virtual .avi stack. These phase contrast image stacks were evaluated after ‘find edges’ processing on ImageJ with further processing using Trackmate 7 plug-in (84) to trace and quantify individual cellular motility tracks (e.g. X, Y positions over time for each cell). Velocity (μm/min) and accumulated distance (μm from start of tracing) data using 0.61μm/pixel as per vendor data (Incucyte Zoom) was further evaluated from exported Trackmate 7 data for each track and additionally represented by motility plots using ImageJ Chemotaxis and Migration Tool plug-in.(85)

### Cell culture confluence and doubling time determination

High resolution phase contrast and brightfield imaging of iPSCs in culture was performed daily for 5 days using the Thrive CellAssist plate imaging instrument (Thrive Biosciences Inc, Beverly MA). Cells were TRA-1-60 positive magnetic sorted (Miltenyi 130-100-832) prior to plating, at 50k/6-well plate well seeding density, 3 wells/line, imaged daily at 4x magnification. Confluence, colony size and doubling times were determined using the CellAssist EvalCore computer workstation

**Figure S1.**
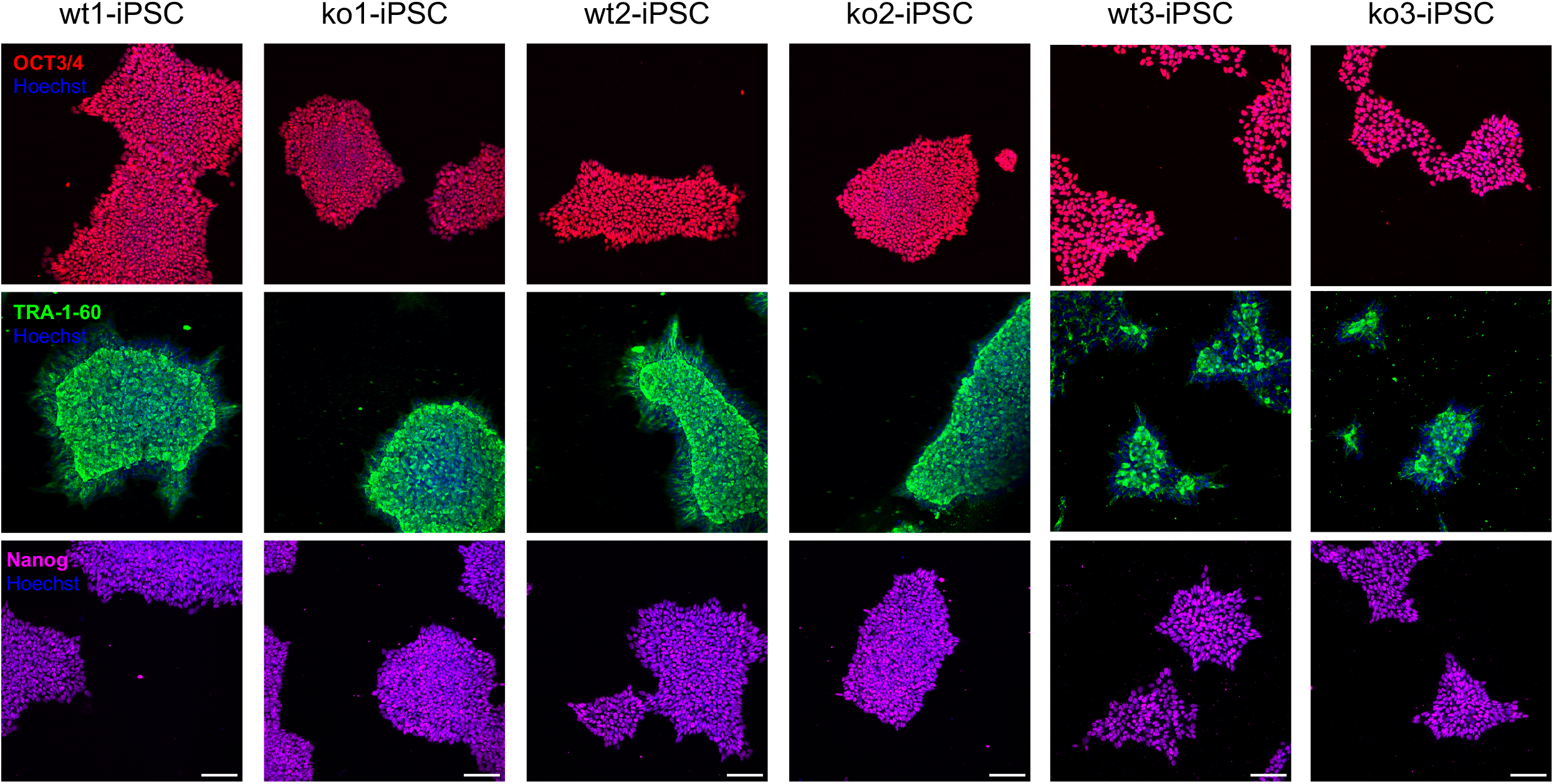
Pluripotency marker immunocytochemistry confirmation of indicated iPSC clones with OCT3/4, TRA-1-60 and Nanog as indicated. Scale bar = 100um

**Figure S2.**
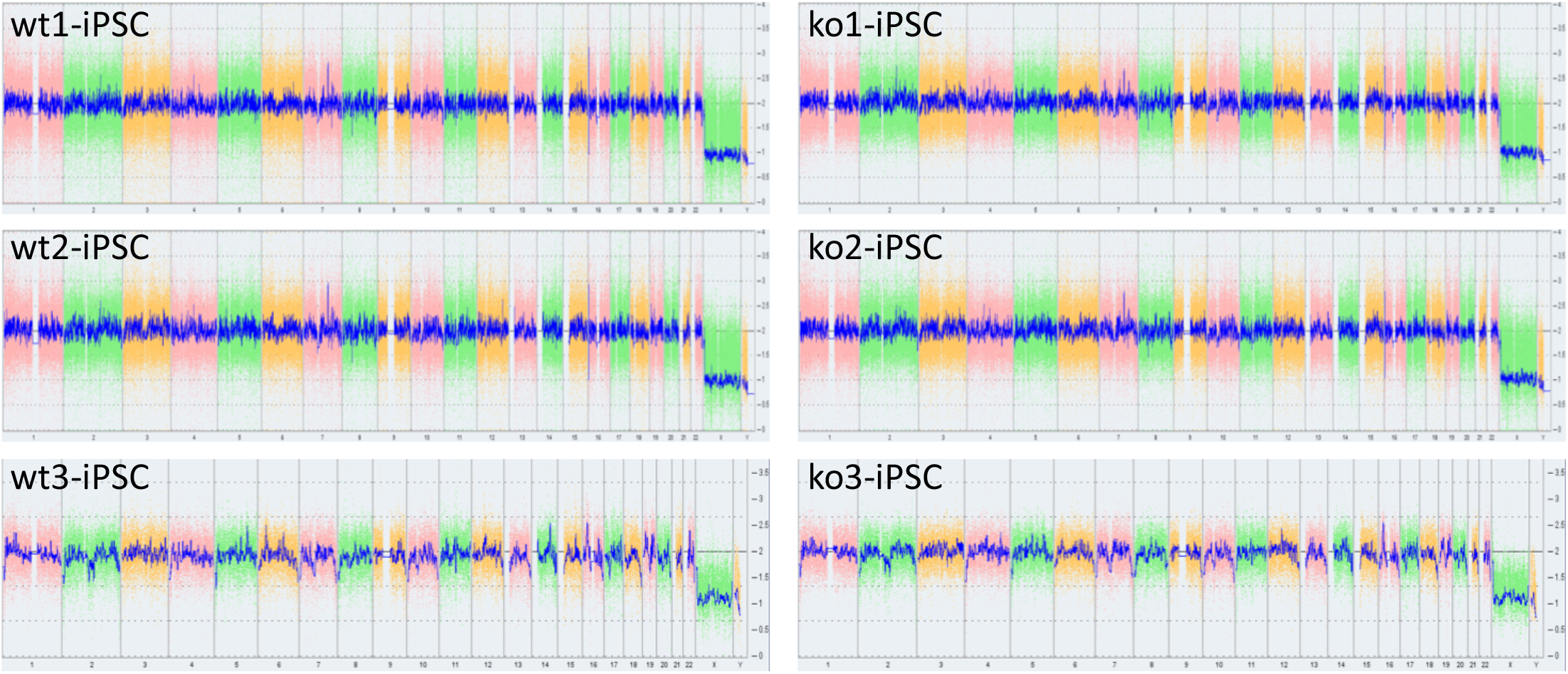
iPSC Karyotypes using Applied Biosystems KaryoStat microarray assay. The whole genome view displays all somatic and sex chromosomes in one frame. The smooth signal plot (right y-axis) is log2 ratios of signal intensities of microarray probes. Values represent normal copy number state (CN = 2), chromosomal gain (CN = 3) or chromosomal loss (CN = 1). The pink, green and yellow colors indicate the raw signal for each individual chromosome probe, while the blue signal represents the normalized probe signal used to determine copy number.

**Figure S3.**
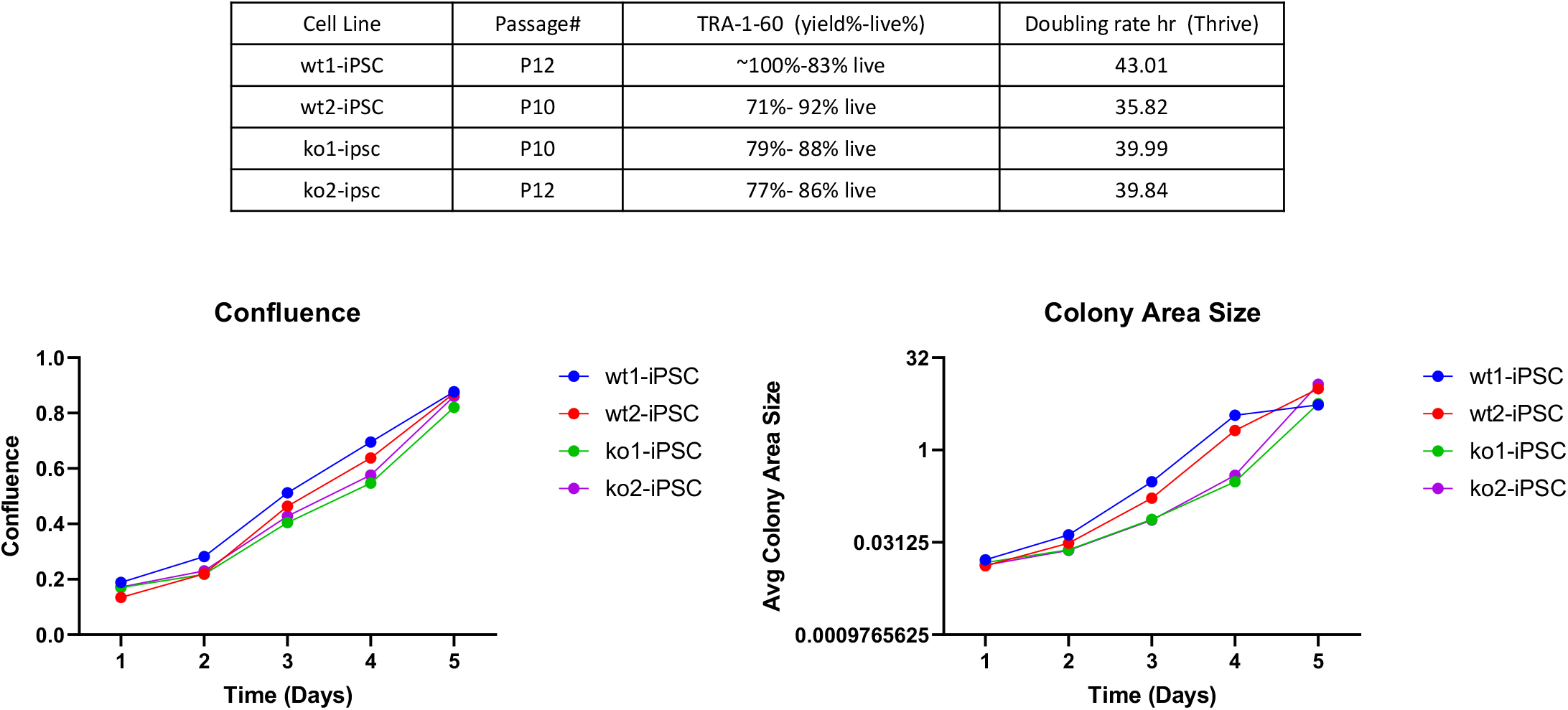
Derivation of CRISPR generated null human iPSC indicate *CYFIP1* expression not required for cellular proliferation or viability. Proliferation doubling time, confluence and colony size over time for indicated WT and KO iPSC lines.

**Figure S4.**
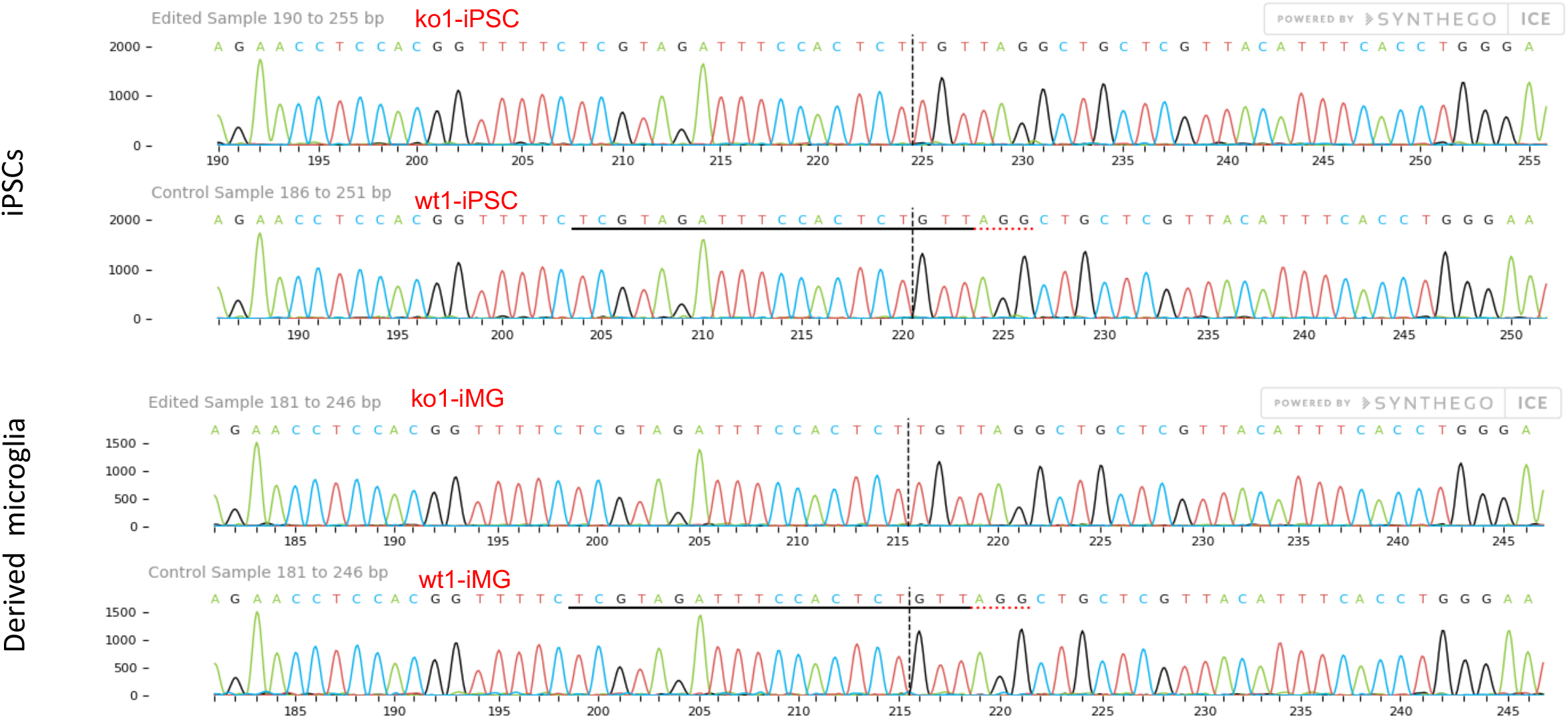
Genotyping of CRISPR *CYFIP1* KO iPSC-MGs (ko1 and wt1-iPSC and iMG lines) confirms retention of loss-of-function upon differentiation.

**Figure S5.**
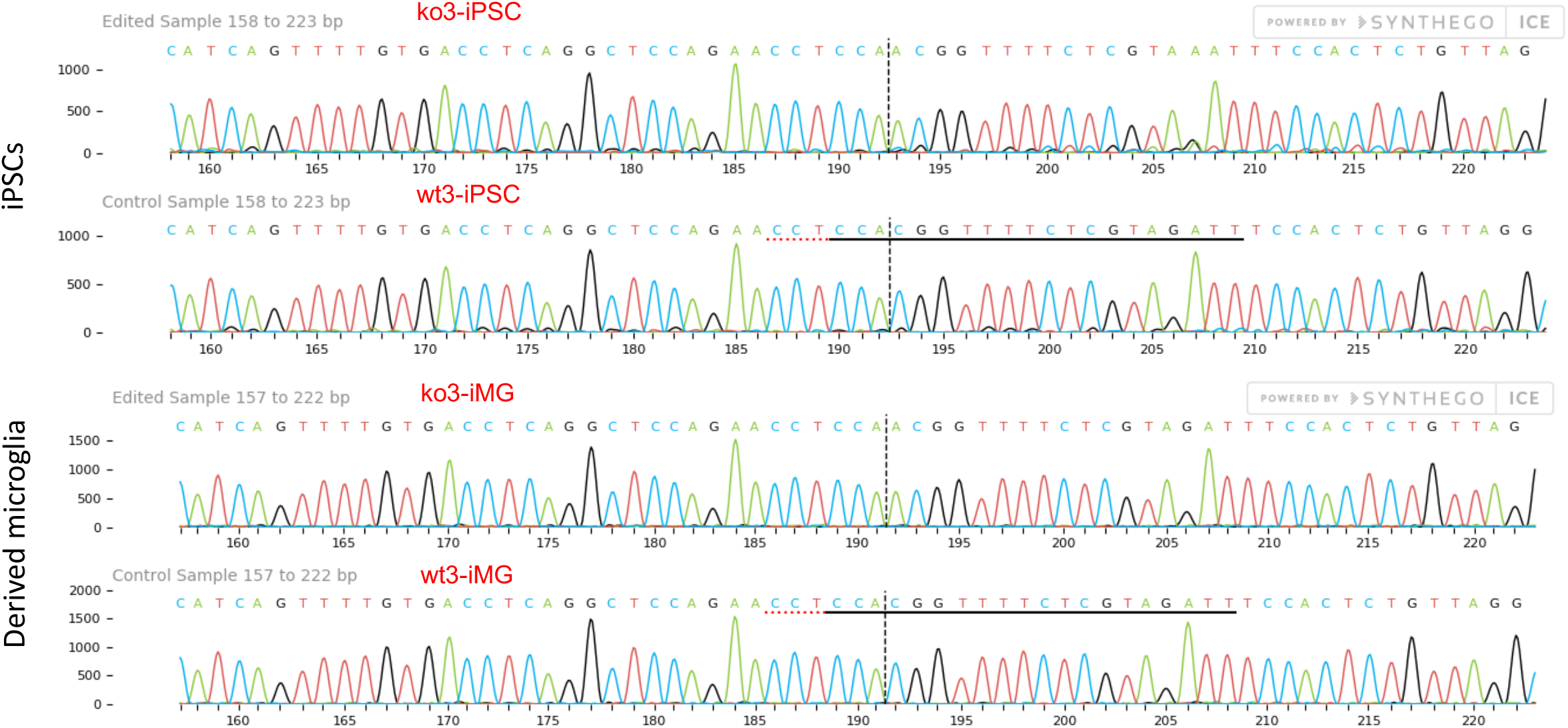
Genotyping of CRISPR CYFIP1 KO iPSC-MGs (ko3 and wt3-iPSC and iMG lines) confirms retention of loss-of-function upon differentiation..

**Table S1.**
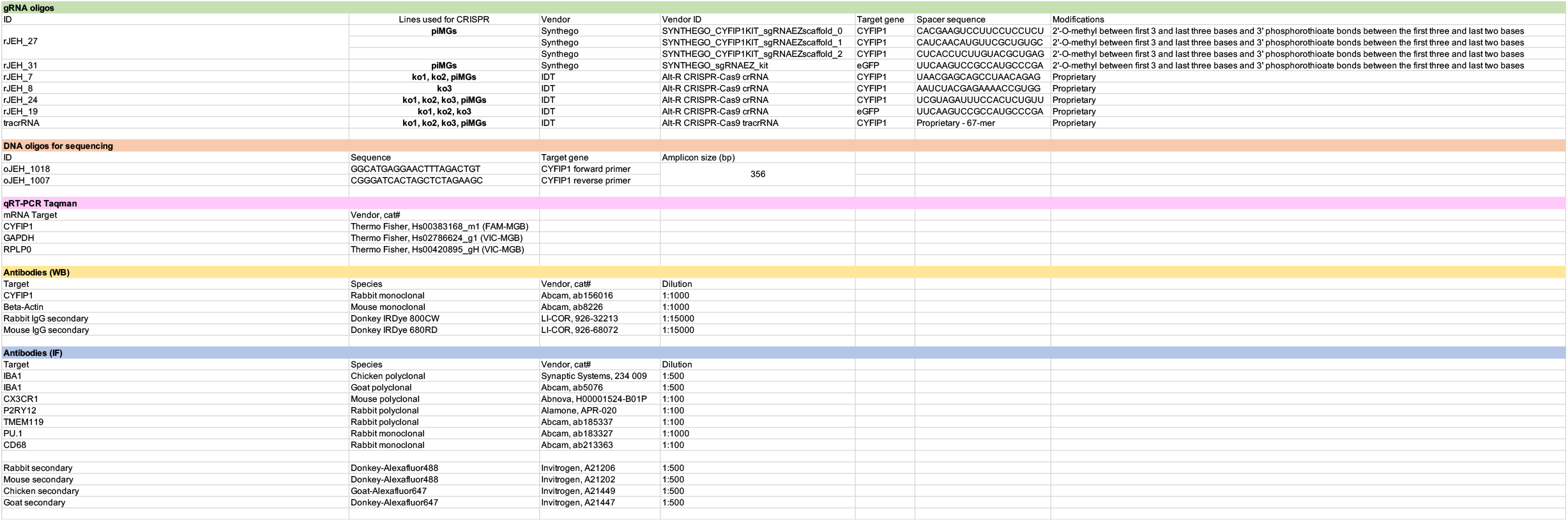
Oligos, gRNAs, primers and antibodies used in this study.

**Table S2.** Cell Lines and assays used in this study

| Parent Line Name | Line Type | Source | Description | Clonal lines use in this study | Assays Included in Study |
| --- | --- | --- | --- | --- | --- |
| VT59122 | PBMC | Vitrologics Inc. | healthy male donor. 40-year old African American | piMGs (not clonal) | Phase contrast, ICC, synaptosome phagocytosis, piMG reprogramming and editing, *CYFIP1* editing sequencing %. |
| PSC-01-179 | iPSC | In-House derivation from fibroblasts | healthy male donor. 29-year old Caucasian, non-Hispanic | wt1-iPSC<br>ko1-iPSC | rtPCR (*CYFIP1*), WB (CYFIP1), *CYFIP1* CRISPR indel locus sequencing, pluripotent marker ICC, karyotype |
| PSC-01-179 | iPSC-MG | In-House derivation from iPSCs | healthy male donor. 29-year old Caucasian, non-Hispanic | wt1-iMG<br>ko1-iMG | synaptosome phagocytosis, rtPCR (*CYFIP1*), WB (CYFIP1), *CYFIP1* CRISPR indel locus sequencing, morphology, motility |
| PSC-01-179 | iPSC | In-House derivation from fibroblasts | healthy male donor. 29-year old Caucasian, non-Hispanic | wt2-iPSC<br>ko2-iPSC | rtPCR (*CYFIP1*), WB (CYFIP1), *CYFIP1* CRISPR indel locus sequencing, pluripotent marker ICC, karyotype |
| PSC-01-179 | iPSC-iMG | In-House derivation from iPSCs | healthy male donor. 29-year old Caucasian, non-Hispanic | wt2-iMG<br>ko2-iMG | synaptosome phagocytosis, rtPCR (*CYFIP1*), iMG ICC marker analysis and quantification, morphology, motility |
| PSC-01-092 | iPSC | In-House derivation from fibroblasts | healthy male donor. 45-year old Caucasian, non-Hispanic | wt3-iPSC<br>ko3-iPSC | rtPCR (*CYFIP1*), WB (CYFIP1), *CYFIP1* CRISPR indel locus sequencing, pluripotent marker ICC, karyotype |
| PSC-01-092 | iPSC-iMG | In-House derivation from iPSCs | healthy male donor. 45-year old Caucasian, non-Hispanic | wt3-iMG,<br>ko3-iMG | synaptosome phagocytosis, motility, *CYFIP1* CRISPR indel locus sequencing |

## References

1. Cox DM, Butler MG (2015): The 15q11. 2 BP1–BP2 microdeletion syndrome: A review. International journal of molecular sciences. 16:4068–4082.

2. Rafi SK, Butler MG (2020): The 15q11. 2 BP1-BP2 Microdeletion (Burnside–Butler) Syndrome: In Silico Analyses of the Four Coding Genes Reveal Functional Associations with Neurodevelopmental Disorders. International Journal of Molecular Sciences. 21:3296.

3. Zhao Q, Li T, Zhao X, Huang K, Wang T, Li Z, et al. (2012): Rare CNVs and Tag SNPs at 15q11.2 Are Associated With Schizophrenia in the Han Chinese Population. Schizophrenia Bulletin. 39:712–719.

4. Rosenfeld JA, Coe BP, Eichler EE, Cuckle H, Shaffer LG (2013): Estimates of penetrance for recurrent pathogenic copy-number variations. Genetics in Medicine. 15:478–481.

5. Vassos E, Collier DA, Holden S, Patch C, Rujescu D, St Clair D, et al. (2010): Penetrance for copy number variants associated with schizophrenia. Human molecular genetics. 19:3477–3481.

6. Pathania M, Davenport EC, Muir J, Sheehan DF, Lopez-Domenech G, Kittler JT (2014): The autism and schizophrenia associated gene CYFIP1 is critical for the maintenance of dendritic complexity and the stabilization of mature spines. Transl Psychiatry. 4:e374.

7. De Rubeis S, Pasciuto E, Li KW, Fernández E, Di Marino D, Buzzi A, et al. (2013): CYFIP1 coordinates mRNA translation and cytoskeleton remodeling to ensure proper dendritic spine formation. Neuron. 79:1169–1182.

8. Oguro-Ando A, Rosensweig C, Herman E, Nishimura Y, Werling D, Bill BR, et al. (2015): Increased CYFIP1 dosage alters cellular and dendritic morphology and dysregulates mTOR. Mol Psychiatry. 20:1069–1078.

9. Santini E, Huynh TN, Longo F, Koo SY, Mojica E, D’Andrea L, et al. (2017): Reducing eIF4E-eIF4G interactions restores the balance between protein synthesis and actin dynamics in fragile X syndrome model mice. Science signaling. 10:eaan0665.

10. Zhang Y, Kang HR, Han K (2019): Differential cell-type-expression of CYFIP1 and CYFIP2 in the adult mouse hippocampus. Animal Cells and Systems. 23:380–383.

11. Saunders A, Macosko EZ, Wysoker A, Goldman M, Krienen FM, de Rivera H, et al. (2018): Molecular Diversity and Specializations among the Cells of the Adult Mouse Brain. Cell. 174:1015-1030.e1016.

12. Zeisel A, Hochgerner H, Lönnerberg P, Johnsson A, Memic F, van der Zwan J, et al. (2018): Molecular Architecture of the Mouse Nervous System. Cell. 174:999-1014.e1022.

13. (2022): CYFIP1 RNA single cell type specificity. Human Protein Atlas proteinatlasorg.

14. Bozdagi O, Sakurai T, Dorr N, Pilorge M, Takahashi N, Buxbaum JD (2012): Haploinsufficiency of Cyfip1 produces fragile X-like phenotypes in mice.

15. Haenseler W, Sansom SN, Buchrieser J, Newey SE, Moore CS, Nicholls FJ, et al. (2017): A Highly Efficient Human Pluripotent Stem Cell Microglia Model Displays a Neuronal-Co-culture-Specific Expression Profile and Inflammatory Response. Stem Cell Reports. 8:1727–1742.

16. Petanjek Z, Judaš M, Šimić G, Rašin MR, Uylings HB, Rakic P, et al. (2011): Extraordinary neoteny of synaptic spines in the human prefrontal cortex. Proceedings of the National Academy of Sciences. 108:13281–13286.

17. Huttenlocher PR (1979): Synaptic density in human frontal cortex-developmental changes and effects of aging. Brain Res. 163:195–205.

18. Faust TE, Gunner G, Schafer DP (2021): Mechanisms governing activity-dependent synaptic pruning in the developing mammalian CNS. Nature Reviews Neuroscience.1-17.

19. Riccomagno MM, Kolodkin AL (2015): Sculpting neural circuits by axon and dendrite pruning. Annu Rev Cell Dev Biol. 31:779–805.

20. Feinberg I (1982): Schizophrenia: Caused by a fault in programmed synaptic elimination during adolescence? Journal of Psychiatric Research. 17:319–334.

21. Sekar A, Bialas AR, de Rivera H, Davis A, Hammond TR, Kamitaki N, et al. (2016): Schizophrenia risk from complex variation of complement component 4. Nature. 530:177–183.

22. Sellgren CM, Gracias J, Watmuff B, Biag JD, Thanos JM, Whittredge PB, et al. (2019): Increased synapse elimination by microglia in schizophrenia patient-derived models of synaptic pruning. Nat Neurosci. 22:374–385.

23. Tam Gloria WC, van de Lagemaat Louie N, Redon R, Strathdee Karen E, Croning Mike DR, Malloy Mary P, et al. (2010): Confirmed rare copy number variants implicate novel genes in schizophrenia. Biochemical Society Transactions. 38:445–451.

24. Yoon K-J, Nguyen HN, Ursini G, Zhang F, Kim N-S, Wen Z, et al. (2014): Modeling a genetic risk for schizophrenia in iPSCs and mice reveals neural stem cell deficits associated with adherens junctions and polarity. Cell stem cell. 15:79–91.

25. Hammond TR, Dufort C, Dissing-Olesen L, Giera S, Young A, Wysoker A, et al. (2019): Single-cell RNA sequencing of microglia throughout the mouse lifespan and in the injured brain reveals complex cell-state changes. Immunity. 50:253-271. e256.

26. Sellgren C, Sheridan S, Gracias J, Xuan D, Fu T, Perlis R (2017): Patient-specific models of microglia-mediated engulfment of synapses and neural progenitors. Molecular Psychiatry. 22.

27. Petazzi P, Menendez P, Sevilla A (2020): CRISPR/Cas9-Mediated Gene Knockout and Knockin Human iPSCs. Methods Mol Biol.

28. Castano J, Bueno C, Jimenez-Delgado S, Roca-Ho H, Fraga MF, Fernandez AF, et al. (2017): Generation and characterization of a human iPSC cell line expressing inducible Cas9 in the “safe harbor” AAVS1 locus. Stem Cell Res. 21:137–140.

29. Ben Jehuda R, Shemer Y, Binah O (2018): Genome Editing in Induced Pluripotent Stem Cells using CRISPR/Cas9. Stem Cell Rev Rep. 14:323–336.

30. Hultquist JF, Schumann K, Woo JM, Manganaro L, McGregor MJ, Doudna J, et al. (2016): A Cas9 Ribonucleoprotein Platform for Functional Genetic Studies of HIV-Host Interactions in Primary Human T Cells. Cell Rep. 17:1438–1452.

31. Schumann K, Lin S, Boyer E, Simeonov DR, Subramaniam M, Gate RE, et al. (2015): Generation of knock-in primary human T cells using Cas9 ribonucleoproteins. Proc Natl Acad Sci U S A. 112:10437–10442.

32. Sheridan SD, Horng JE, Perlis RH (2022): Patient-derived in vitro models of microglial function and synaptic engulfment in schizophrenia. Biological Psychiatry.

33. Ginhoux F, Greter M, Leboeuf M, Nandi S, See P, Gokhan S, et al. (2010): Fate mapping analysis reveals that adult microglia derive from primitive macrophages. Science. 330:841–845.

34. Schulz C, Gomez Perdiguero E, Chorro L, Szabo-Rogers H, Cagnard N, Kierdorf K, et al. (2012): A lineage of myeloid cells independent of Myb and hematopoietic stem cells. Science. 336:86–90.

35. Hoeffel G, Ginhoux F (2018): Fetal monocytes and the origins of tissue-resident macrophages. Cell Immunol. 330:5–15.

36. Abud EM, Ramirez RN, Martinez ES, Healy LM, Nguyen CHH, Newman SA, et al. (2017): iPSC-Derived Human Microglia-like Cells to Study Neurological Diseases. Neuron. 94:278–293 e279.

37. Banerjee P, Paza E, Perkins EM, James OG, Kenkhuis B, Lloyd AF, et al. (2020): Generation of pure monocultures of human microglia-like cells from induced pluripotent stem cells. Stem Cell Research. 49:102046.

38. Douvaras P, Sun B, Wang M, Kruglikov I, Lallos G, Zimmer M, et al. (2017): Directed differentiation of human pluripotent stem cells to microglia. Stem cell reports. 8:1516–1524.

39. McQuade A, Coburn M, Tu CH, Hasselmann J, Davtyan H, Blurton-Jones M (2018): Development and validation of a simplified method to generate human microglia from pluripotent stem cells. Mol Neurodegener. 13:67.

40. Campellone KG, Welch MD (2010): A nucleator arms race: cellular control of actin assembly. Nature Reviews Molecular Cell Biology. 11:237–251.

41. Rotty JD, Wu C, Bear JE (2013): New insights into the regulation and cellular functions of the ARP2/3 complex. Nature Reviews Molecular Cell Biology. 14:7–12.

42. Gautreau AM, Fregoso FE, Simanov G, Dominguez R (2022): Nucleation, stabilization, and disassembly of branched actin networks. Trends in Cell Biology. 32:421–432.

43. Drew J, Arancibia-Carcamo IL, Jolivet R, Lopez-Domenech G, Attwell D, Kittler JT (2020): Control of microglial dynamics by Arp2/3 and the autism and schizophrenia-associated protein Cyfip1. bioRxiv.

44. Lynch MA (2009): The multifaceted profile of activated microglia. Molecular neurobiology. 40:139–156.

45. Stirling DR, Swain-Bowden MJ, Lucas AM, Carpenter AE, Cimini BA, Goodman A (2021): CellProfiler 4: improvements in speed, utility and usability. BMC Bioinformatics. 22:433.

46. Nimmerjahn A, Kirchhoff F, Helmchen F (2005): Resting Microglial Cells Are Highly Dynamic Surveillants of Brain Parenchyma in Vivo. Science. 308:1314–1318.

47. Verney C, Monier A, Fallet-Bianco C, Gressens P (2010): Early microglial colonization of the human forebrain and possible involvement in periventricular white-matter injury of preterm infants. Journal of Anatomy. 217:436–448.

48. Monier A, Evrard P, Gressens P, Verney C (2006): Distribution and differentiation of microglia in the human encephalon during the first two trimesters of gestation. Journal of Comparative Neurology. 499:565–582.

49. Mosser C-A, Baptista S, Arnoux I, Audinat E (2017): Microglia in CNS development: Shaping the brain for the future. Progress in neurobiology. 149:1–20.

50. Cunningham CL, Martínez-Cerdeño V, Noctor SC (2013): Microglia regulate the number of neural precursor cells in the developing cerebral cortex. Journal of Neuroscience. 33:4216–4233.

51. Diaz-Aparicio I, Paris I, Sierra-Torre V, Plaza-Zabala A, Rodríguez-Iglesias N, Márquez- Ropero M, et al. (2020): Microglia Actively Remodel Adult Hippocampal Neurogenesis through the Phagocytosis Secretome. The Journal of Neuroscience. 40:1453.

52. Sierra A, Encinas JM, Deudero JJP, Chancey JH, Enikolopov G, Overstreet-Wadiche LS, et al. (2010): Microglia Shape Adult Hippocampal Neurogenesis through Apoptosis-Coupled Phagocytosis. Cell Stem Cell. 7:483–495.

53. Sato K (2015): Effects of microglia on neurogenesis. Glia. 63:1394–1405.

54. Pérez-Rodríguez DR, Blanco-Luquin I, Mendioroz M (2021): The participation of microglia in neurogenesis: a review. Brain sciences. 11:658.

55. Tremblay M-È, Lowery RL, Majewska AK (2010): Microglial interactions with synapses are modulated by visual experience. PLoS biology. 8:e1000527.

56. Paolicelli RC, Bolasco G, Pagani F, Maggi L, Scianni M, Panzanelli P, et al. (2011): Synaptic pruning by microglia is necessary for normal brain development. Science. 333:1456–1458.

57. Glausier JR, Lewis DA (2013): Dendritic spine pathology in schizophrenia. Neuroscience. 251:90–107.

58. Konopaske GT, Lange N, Coyle JT, Benes FM (2014): Prefrontal cortical dendritic spine pathology in schizophrenia and bipolar disorder. JAMA Psychiatry. 71:1323–1331.

59. Neniskyte U, Gross CT (2017): Errant gardeners: glial-cell-dependent synaptic pruning and neurodevelopmental disorders. Nature Reviews Neuroscience. 18:658–670.

60. Stevens B, Allen NJ, Vazquez LE, Howell GR, Christopherson KS, Nouri N, et al. (2007): The classical complement cascade mediates CNS synapse elimination. Cell. 131:1164–1178.

61. Schafer DP, Lehrman EK, Kautzman AG, Koyama R, Mardinly AR, Yamasaki R, et al. (2012): Microglia sculpt postnatal neural circuits in an activity and complement-dependent manner. Neuron. 74:691–705.

62. Paolicelli RC, Bolasco G, Pagani F, Maggi L, Scianni M, Panzanelli P, et al. (2011): Synaptic pruning by microglia is necessary for normal brain development. Science. 333:1456–1458.

63. Aguzzi A, Barres BA, Bennett ML (2013): Microglia: scapegoat, saboteur, or something else? Science. 339:156–161.

64. Miyanishi K, Sato A, Kihara N, Utsunomiya R, Tanaka J (2021): Synaptic elimination by microglia and disturbed higher brain functions. Neurochemistry International. 142:104901.

65. Waltes R, Duketis E, Knapp M, Anney RJ, Huguet G, Schlitt S, et al. (2014): Common variants in genes of the postsynaptic FMRP signalling pathway are risk factors for autism spectrum disorders. Human genetics. 133:781–792.

66. Wang J, Tao Y, Song F, Sun Y, Ott J, Saffen D (2015): Common regulatory variants of CYFIP1 contribute to susceptibility for autism spectrum disorder (ASD) and classical autism. Annals of human genetics. 79:329–340.

67. Augusto-Oliveira M, Arrifano GP, Delage CI, Tremblay MÈ, Crespo-Lopez ME, Verkhratsky A (2022): Plasticity of microglia. Biological Reviews. 97:217–250.

68. Savage JC, Carrier M, Tremblay M-È (2019): Morphology of Microglia Across Contexts of Health and Disease. In: Garaschuk O, Verkhratsky A, editors. Microglia: Methods and Protocols. New York, NY: Springer New York, pp 13-26.

69. Donat CK, Scott G, Gentleman SM, Sastre M (2017): Microglial activation in traumatic brain injury. Frontiers in aging neuroscience. 9:208.

70. Au NPB, Ma CHE (2017): Recent advances in the study of bipolar/rod-shaped microglia and their roles in neurodegeneration. Frontiers in aging neuroscience. 9:128.

71. Graeber MB (2010): Changing face of microglia. Science. 330:783–788.

72. Streit WJ, Sammons NW, Kuhns AJ, Sparks DL (2004): Dystrophic microglia in the aging human brain. Glia. 45:208–212.

73. Karperien A, Ahammer H, Jelinek HF (2013): Quantitating the subtleties of microglial morphology with fractal analysis. Frontiers in cellular neuroscience. 7:3.

74. Colombo G, Cubero RJA, Kanari L, Venturino A, Schulz R, Scolamiero M, et al. (2022): A tool for mapping microglial morphology, morphOMICs, reveals brain-region and sex-dependent phenotypes. Nature Neuroscience. 1–15.

75. Smolders SM-T, Kessels S, Vangansewinkel T, Rigo J-M, Legendre P, Brône B (2019): Microglia: Brain cells on the move. Progress in Neurobiology. 178:101612.

76. Hattori Y, Naito Y, Tsugawa Y, Nonaka S, Wake H, Nagasawa T, et al. (2020): Transient microglial absence assists postmigratory cortical neurons in proper differentiation. Nature Communications. 11:1631.

77. Sellgren CM, Sheridan SD, Gracias J, Xuan D, Fu T, Perlis RH (2017): Patient-specific models of microglia-mediated engulfment of synapses and neural progenitors. Mol Psychiatry. 22:170–177.

78. Sheridan SD, Thanos JM, De Guzman RM, McCrea LT, Horng JE, Fu T, et al. (2021): Umbilical cord blood-derived microglia-like cells to model COVID-19 exposure. Translational psychiatry. 11:1–9.

79. Gray EG, Whittaker VP (1962): The isolation of nerve endings from brain: an electron-microscopic study of cell fragments derived by homogenization and centrifugation. J Anat. 96:79–88.

80. Kamat PK, Kalani A, Tyagi N (2014): Method and validation of synaptosomal preparation for isolation of synaptic membrane proteins from rat brain. MethodsX. 1:102–107.

81. Tenreiro P, Rebelo S, Martins F, Santos M, Coelho ED, Almeida M, et al. (2017): Comparison of simple sucrose and percoll based methodologies for synaptosome enrichment. Anal Biochem. 517:1–8.

82. Schindelin J, Arganda-Carreras I, Frise E, Kaynig V, Longair M, Pietzsch T, et al. (2012): Fiji: an open-source platform for biological-image analysis. Nature Methods. 9:676–682.

83. Li K (2008): The image stabilizer plugin for ImageJ.

84. Ershov D, Phan M-S, Pylvänäinen JW, Rigaud SU, Le Blanc L, Charles-Orszag A, et al. (2022): TrackMate 7: integrating state-of-the-art segmentation algorithms into tracking pipelines. Nature Methods. 19:829–832.

85. Zantl R, Horn E (2011): Chemotaxis of Slow Migrating Mammalian Cells Analysed by Video Microscopy. In: Wells CM, Parsons M, editors. Cell Migration: Developmental Methods and Protocols. Totowa, NJ: Humana Press, pp 191–203.

